# Identification of protein aggregates in the aging vertebrate brain with prion-like and phase separation properties

**DOI:** 10.1101/2022.02.26.482115

**Authors:** Itamar Harel, Yiwen R. Chen, Inbal Ziv, Param Priya Singh, Paloma Navarro Negredo, Uri Goshtchevsky, Wei Wang, Gwendoline Astre, Eitan Moses, Andrew McKay, Ben E. Machado, Katja Hebestreit, Sifei Yin, Alejandro Sánchez Alvarado, Daniel F. Jarosz, Anne Brunet

**Author notes:** These authors contributed equally (Itamar Harel, Yiwen R. Chen). These authors contributed equally (Inbal Ziv, Param Priya Singh). Senior authors (Daniel F. Jarosz, Anne Brunet).

## Abstract

Protein aggregation, which can sometimes spread in a prion-like manner, is a hallmark of neurodegenerative diseases. However, whether prion-like aggregates form during normal brain aging remains unknown. Here we use quantitative proteomics in the African turquoise killifish to identify protein aggregates that accumulate in old vertebrate brains. These aggregates are enriched for prion-like RNA binding proteins, notably the ATP-dependent RNA helicase DDX5. We validate that DDX5 forms mislocalized cytoplasmic aggregates in the brains of old killifish and mice. Interestingly, DDX5’s prion-like domain allows these aggregates to propagate across many generations in yeast. *In vitro*, DDX5 phase separates into condensates. Mutations that abolish DDX5 prion propagation also impair the protein’s ability to phase separate. DDX5 condensates exhibit enhanced enzymatic activity, but they can mature into inactive, solid aggregates. Our findings suggest that protein aggregates with prion-like properties form during normal brain aging, which could have implications for the age-dependency of cognitive decline.

## INTRODUCTION

Aging is accompanied by a striking increase in the incidence of neurodegenerative diseases. Many of these diseases are characterized by progressive accumulation of protein aggregates in the brain (e.g., Amyloid-β and Tau for Alzheimer’s disease, α-Synuclein for Parkinson’s disease, and Huntingtin for Huntington’s disease) (Dugger and Dickson, 2017; Forman et al., 2004). Protein aggregates have also been shown to accumulate during normal aging in several non-vertebrate organisms, such as *C. elegans* and yeast (Ben-Zvi et al., 2009; David et al., 2010; Huang et al., 2019; Labbadia and Morimoto, 2014; Lechler et al., 2017; Lopez-Otin et al., 2013; Walther et al., 2015; Wolff et al., 2014). While the aging proteome has started to be characterized in vertebrates in various tissues, including the brain (Angelidis et al., 2019; Gebert et al., 2020; Kelmer Sacramento et al., 2020; Ori et al., 2015; Xiao et al., 2020; Yu et al., 2020), a systematic examination of protein aggregates during aging is missing.

Prion-like properties have been proposed to drive the progression of several neurodegenerative pathologies by facilitating transmission of protein aggregates from affected to unaffected areas of the brain (Brundin and Melki, 2017; Brundin et al., 2010; Collinge, 2016; Laferriere et al., 2019; Walker and Jucker, 2015). Long-viewed as a rare biological oddity, prions have recently been discovered throughout evolution, from yeast to humans (Halfmann, 2016; Halfmann et al., 2010; Harvey et al., 2017). Prions and prion-like self-assembly have also been implicated in multiple normal physiological functions, such as metabolism, cell fate determination, antiviral responses, and inflammation (Cai et al., 2014; Chakrabortee et al., 2016; Garcia et al., 2016; Halfmann, 2016; Harvey et al., 2017; Hervas et al., 2020; Holmes et al., 2013; Jarosz and Khurana, 2017). Although proteins with prion-like properties were identified in old *C. elegans* (Huang et al., 2019), whether prion-like protein aggregates accumulate during vertebrate aging, and notably in the brain, is unknown.

The African turquoise killifish *Nothobranchius furzeri* is an ideal vertebrate model to investigate protein aggregation in the aging brain. Its lifespan is 6-10 times shorter than mice or zebrafish (Cellerino et al., 2016; Valdesalici and Cellerino, 2003). Within this short time frame, the killifish recapitulates defining features of vertebrate aging: cognitive decline, decline in fertility, sarcopenia, and neurofibrillary degeneration in the brain (Harel and Brunet, 2015; Kim et al., 2016; Matsui et al., 2019; Terzibasi et al., 2007; Valenzano et al., 2006a; Valenzano et al., 2006b). Moreover, the killifish responds to conserved lifespan interventions, including dietary restriction (Terzibasi et al., 2009; Valenzano et al., 2006b), and integrative genomic and genome-editing toolkits have been developed in the organism (Harel et al., 2015; Harel et al., 2016; Reichwald et al., 2015; Valenzano et al., 2015; Valenzano et al., 2011). Some protein aggregates have been examined qualitatively in aged killifish brains, such as ribosome subunits (Kelmer Sacramento et al., 2020). However, systematic and quantitative identification of proteins that aggregate in old brains, and whether they have a prion-like behavior, has never been done.

Here we leverage the African killifish as a powerful model to unbiasedly identify proteins that aggregate during normal brain aging. Using quantitative proteomics, we identify many proteins with an increased propensity to aggregate in the aging brain. One of these proteins, the RNA helicase DDX5, forms mislocalized cytoplasmic aggregates in old brains of both killifish and mice. We show that DDX5 has prion-like seeding properties both *in vitro* and in a yeast system. *In vitro*, DDX5 rapidly undergoes phase transition under physiological conditions, which may underlie its seeding properties. Although DDX5 is highly active in its droplet form, these condensates mature into solid aggregates that are inactive and potentially infectious. Collectively, our results show that many proteins aggregate during normal vertebrate brain aging and raise the possibility that these aggregates could contribute to the age-dependency of cognitive decline.

## RESULTS

### Proteomic identification of proteins that aggregate in the old vertebrate brain reveals enrichment for prion-like domains

The identity of protein aggregates that arise during normal brain aging is poorly understood. To identify such aggregates, we used a protocol that separates them from large protein complexes and membraneless organelles based on molecular weight. This approach was developed to detect amyloids (Kryndushkin et al., 2013) and identify other types of protein aggregates (Chen, Harel, et al.). Briefly, we collected the brains of three young (3.5 months) and three old (7 months) male killifish (Figure 1A; we also collected six other tissues from these fish and old telomerase (TERT)-mutant fish (described in Chen, Harel, et al.). Using a series of differential centrifugations, we isolated a fraction enriched for high molecular weight protein aggregates (AGG) and another fraction of matched native tissue lysate (TL) (see Chen, Harel, et al.). The aged brain aggregate fraction displayed substantially more SDS-resistant species (Figure 1B, inset, see also Chen, Harel, et al.), consistent with the notion that protein aggregation is enhanced in the killifish brain during normal aging.

**Figure 1:**
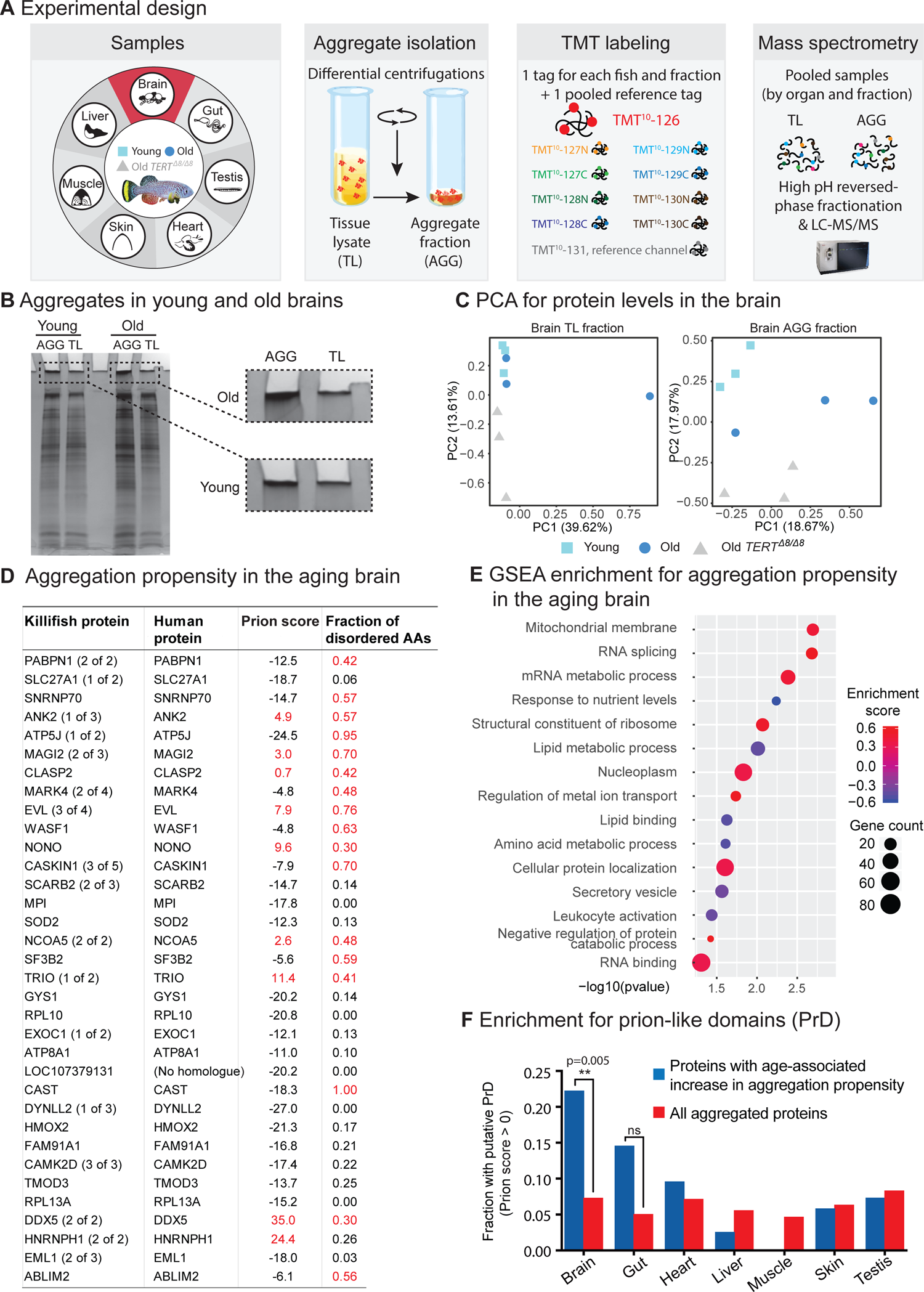
Systematic and quantitative identification of age-related protein aggregation in the vertebrate brain. **(A)** Quantitative proteomic analysis pipeline for the brain (and other organs) from young, old, and old *TERT^Δ8/Δ8^*male fish. For each organ, a tissue lysate (TL) and an aggregate fraction (AGG) were isolated, and tandem mass tag (TMT) labeling and mass spectrometry was performed. **(B)** Silver stain of SDS-PAGE gel with TL and AGG samples (treated with 2% SDS without boiling) from young and old brains highlights the presence of aggregates in the stacking gel. Equal amounts (1 µg) of tissue lysate (TL) and aggregate (AGG) fraction from brain and liver were used for silver stain analysis. A version of the gel with additional organs is shown in Figure 1C of Chen, Harel, et al. **(C)** Principal component analysis (PCA) on protein abundance in either the tissue lysate (TL) or the aggregated (AGG) fractions from the brain of young (light blue square), old (dark blue circle), and old *TERT^Δ8/Δ8^*(grey triangles) fish. Each symbol represents an individual fish. Another version of this brain PCA is included with that of other tissues in Figure 1E of Chen, Harel, et al. **(D)** Ranked list of proteins with increased aggregation propensity in old brains compared to young brains. Ranking is based on descending fold change for aggregation propensity during aging – i.e. the relative abundance ratio of aggregate versus tissue lysate, averaged across three fish in old versus young killifish brain (p<0.05, Student’s t-test). Range is 3.5 to 1.5 fold. Predicted prion-like domain (PrD) scores (using PLAAC) and fraction of disordered amino acids (AAs) (using DISOPRED3) are presented for each protein. Red: PrD score > 0 and fraction of disordered AAs > 0.3 (30%) are considered to be prion-like proteins or intrinsically disordered proteins, respectively. Paralogs are indicated “X of Y” (e.g., “2 of 3” means the second paralog out of three identified paralogs). See also Table S2 in Chen, Harel, et al. **(E)** GO enrichment using Gene Set Enrichment Analysis (GSEA) for proteins with an increase in aggregation propensity in old brains. Enrichment score = score for enrichment/depletion of GO terms in this gene set. Counts = number of genes. **(F)** Enrichment for prion-like domains (PrD) in proteins with increased aggregation propensity with age in the brain and other tissues. Fraction with putative PrD (PrD score > 0). Red: all detected aggregated proteins; Blue: proteins with increased aggregation propensity with aging (t-value >=0.75, p<0.05 from Student’s t-test). Age-associated differences in aggregation propensity was assessed using Fisher’s exact test.

Next, we quantitatively characterized protein aggregation in old brains using tandem mass tag (TMT)-based quantitative Mass Spectrometry (MS) (Figure 1A) (see Chen, Harel, et al.). Principal Component Analysis (PCA) revealed stronger separation on the aggregate fractions than on total lysate (Figure 1C). To identify proteins that are intrinsically more prone to aggregation with age, we calculated the aggregation propensity, the relative abundance ratio of aggregate versus tissue lysate, for each protein expressed in the brain (Figure 1D, list ranked in descending order from 3.5 to 1.5 fold). Proteins with significant age-dependent increases in aggregation propensity in the brain included many proteins with RNA binding properties (Cai et al., 2014; Chakrabortee et al., 2016; Garcia et al., 2016; Gebauer et al., 2021; Halfmann, 2016; Harvey et al., 2017; Hervas et al., 2020; Holmes et al., 2013; Jarosz and Khurana, 2017; Sharma et al., 2021), including proteins involved in splicing (Figures 1D and E, Figure S1A). Thus, this proteomic dataset identifies many candidate proteins with an increased propensity to aggregate in old killifish brains.

Interestingly, brain proteins with an increased aggregation propensity with age were uniquely enriched for protein sequences that are predicted to have prion-like properties (Lancaster et al., 2014) (Figures 1D (red) and F). In addition, they were also enriched for the presence of intrinsically disordered regions (Jones and Cozzetto, 2015) – domains that are not predicted to adopt a single three-dimensional structure in solution – another hallmark of prion and prion-like proteins (Jones and Cozzetto, 2015) (Figures 1D and F, Figure S1B). Collectively, these data identify proteins (e.g., RNA binding proteins) that form aggregates in the vertebrate brains during physiological aging and reveal an enrichment for proteins with disordered and prion-like domains.

### The RNA helicase DDX5 forms aggregate-like puncta in old vertebrate brains and when overexpressed

Among the proteins that aggregate with age in the killifish brain, we focused on DDX5 – an ATP-dependent RNA helicase – because of its strong prion-like score (Figure 1D) and its role in RNA splicing (Dardenne et al., 2014). DDX5 has a C-terminal intrinsically disordered region (IDR), which encompasses a domain with a strong prion-like score (PrD) (Figures 2A and B, Figures S2A and S2B). Both the IDR entire region and the specific PrD are conserved in human DDX5 (Figures 2A and 2B).

**Figure 2:**
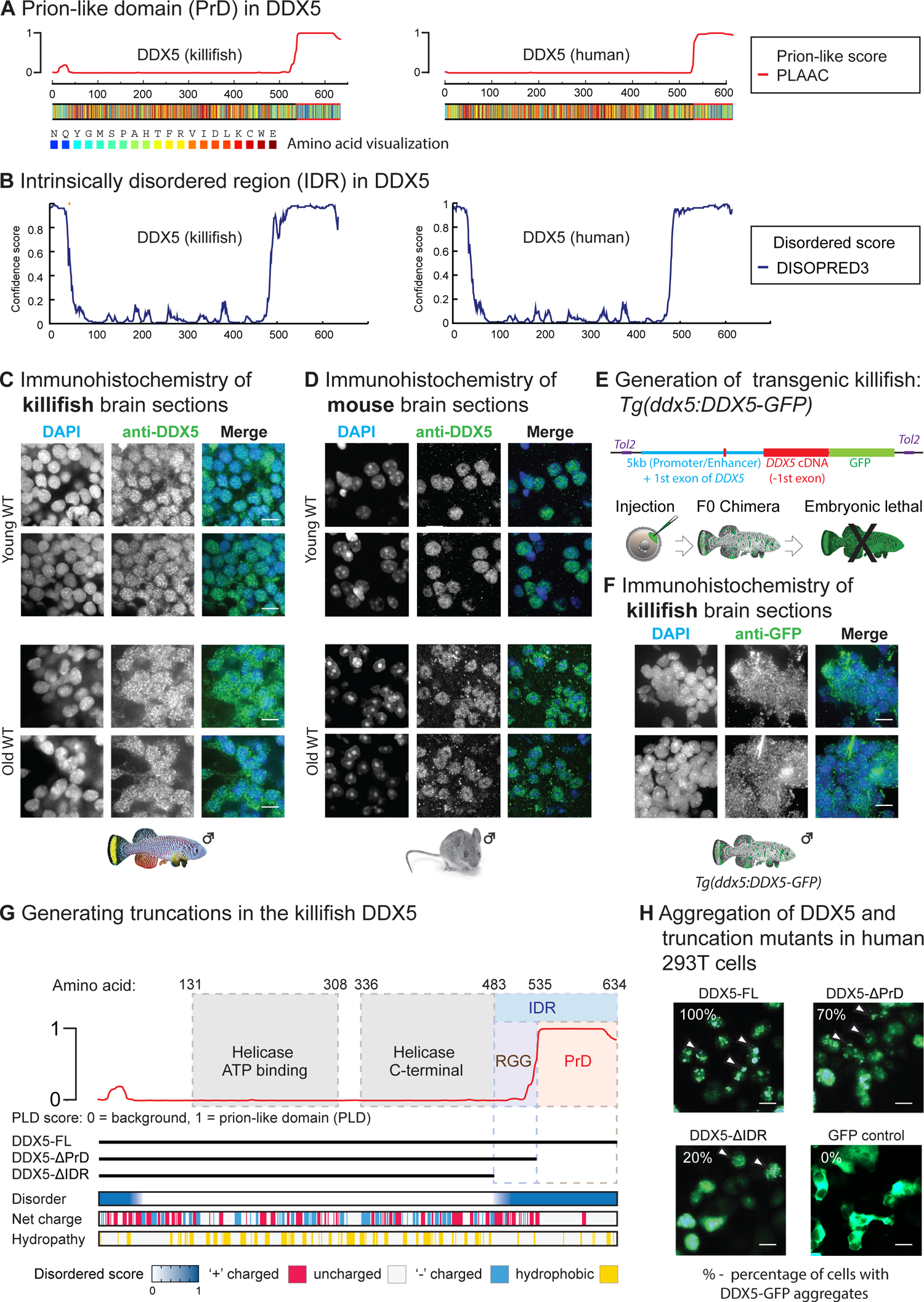
The RNA helicase DDX5 has a conserved prion-like domain and shows increased aggregate-like puncta in old brains in killifish and mammals. **(A)** The presence and location of the predicted prion-like (PrD) domain in the DDX5 protein is conserved between killifish and humans. Red line: PLAAC score of the protein sequence. DDX5 amino acid positions are indicated at the bottom. Amino-acid composition is color-coded for visualization. **(B)** The presence and location of the predicted intrinsically disordered region (IDR) in the DDX5 protein is conserved between killifish and humans. Blue line: predicted amino acid disorder score based on DISOPRED3. **(C)** Immunohistochemistry for DDX5 in brain sections from young (3.5 months) and old (7 months) male killifish. Green: DDX5, Blue: DAPI (nuclei). Image representative of 3 individual fish for each age group. Staining was performed twice independently, with a total of ∼7 sections per fish per experiment. Scale bar: 10µm. Quantification of the subcellular localization of DDX5 puncta is in Figure S2E. **(D)** Immunohistochemistry for DDX5 in brain sections from young (4 months) and old (28 months) male mice. Green: DDX5, Blue: DAPI (nuclei). Image representative of 2 mice per age group, with sections per mouse. Scale bar: 20µm. **(E)** The *Tg(ddx5:DDX5-GFP)* transgenic line was generated by using 5kb surrounding the transcriptional start site of the *DDX5* gene (including the first exon) to drive the expression of DDX5-GFP fusion protein. As *Tg(ddx5:DDX5-GFP)* were not viable as F1 fish, we used F0 chimeric adult fish (∼6-7 months old) for the staining. **(F)** Immunohistochemistry for GFP in brain sections from old (∼6-7 months) male *Tg(ddx5:DDX5-GFP)* transgenic killifish. Green: GFP, Blue: DAPI (nuclei). Image representative of 3 individual fish, ∼6 sections per fish. Scale bar: 10µm. **(G)** Generation of killifish DDX5 mutants with truncation of the prion-like domain (DDX5-ΔPrD) or truncation of the intrinsically disordered region (DDX5-ΔIDR) tagged with EGFP for expression in mammalian cells. FL: full length. Red line: PLAAC score of the protein sequence. Bottom: Color-coded distribution of the disordered region (DISOPRED3, blue), net charge (sliding window of 5 amino acids), and hydropathy (sliding window of 5 amino acids) in the killifish DDX5 protein. **(H)** Percentage of aggregate-like fluorescent puncta of DDX5-EGFP full length (FL) and truncation mutants in human 293T cells. Images are representative of two independent experiments, each performed in triplicate wells. Scale bar: 10µm. The percentage of cells with DDX5-GFP aggregates (estimated based on >3 fields of view for each replicate) is indicated in the top left corner.

To independently test whether DDX5 forms aggregates in the aging killifish brain, we generated an antibody to killifish DDX5 (Figure S2C) and performed immunostaining on brain sections. These immunostaining data revealed that DDX5 is present in nuclear puncta in young brains, but that these puncta become disorganized and mislocalized (more cytoplasmic) in old killifish brains (Figure 2C, Figures S2D and S2E). Interestingly, immunostaining of young and old mouse brains with antibodies to mammalian DDX5 also showed increased disorganized and mislocalized cytoplasmic puncta in the old brain (Figure 2D). Thus, DDX5 assembles in nuclear puncta in young animals, but these puncta become disorganized and mislocalized in old vertebrate brains.

We then asked if exogenous overexpression of DDX5 can result in protein aggregation. To this end, we generated a line of transgenic fish expressing killifish DDX5 fused to GFP driven by *DDX5* regulatory regions (Figure 2E, Figure S3A). We found that overexpressed DDX5-GFP also formed mislocalized cytoplasmic fluorescent puncta in the brain (even already in young) (Figure 2F). We also generated DDX5 knock-out killifish and found that they were not viable, highlighting the importance of this RNA helicase at the organismal level (Figure S3B). When overexpressed in human 293T cells, killifish DDX5 also formed cytoplasmic and nuclear puncta (Figures 2G and H, Figure S3C). Truncation of the entire IDR region of DDX5 abolished the formation of protein aggregates in cultured cells (Figures 2G and H). In contrast, truncation of the prion-like domain (PrD) alone still allowed protein aggregation (albeit less than wild-type) (Figures 2G and H).

Together, these results establish that DDX5, which was identified by mass spectrometry to have an increased propensity to aggregate during aging in killifish brains, also aggregates *in vivo* in old vertebrate brains and in mammalian cells. These findings also identify the disordered domain of DDX5 as a critical molecular determinant of its aggregation capacity.

### DDX5 aggregates act as prions that can seed the formation of new aggregates

We next asked whether DDX5 aggregates have ‘infectious’ prion-like properties – i.e., the ability to nucleate or ‘seed’ the formation of larger protein aggregates in an autocatalytic manner. To this end, we used an inducible system (Weinhandl et al., 2014) in which yeast cells are switched from raffinose to galactose medium to drive the expression of fluorescently-tagged fusion proteins (Figure 3A). A hallmark of prion proteins is that their overexpression induces their aggregation (Wickner, 1994). When overexpressed in yeast, killifish DDX5-EGFP (and other proteins that aggregated in old brains, e.g., MARK4, SCARB2) formed aggregate-like fluorescent puncta, whereas EGFP (or mRuby3) fused to other proteins (e.g., PABPN1 and HNRNPH1) did not (Figures 3B and C).

**Figure 3:**
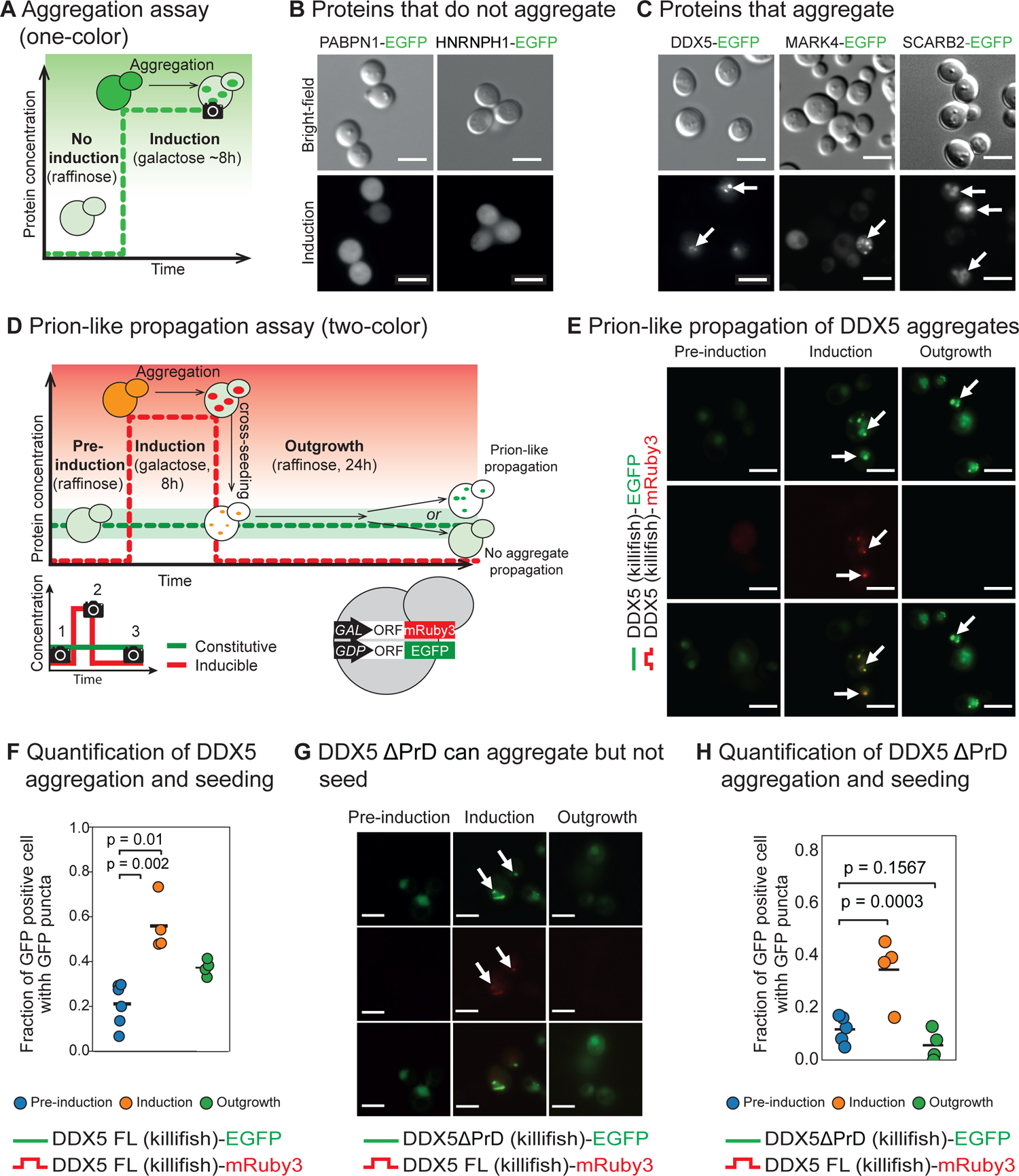
DDX5 forms aggregate-like puncta in yeast and exhibits prion-like seeding properties that are dependent on the PrD domain. **(A)** ‘One-color’ experimental design to assess protein aggregation in yeast, by conditionally expressing C-terminally EGFP-tagged killifish protein candidates under a galactose-inducible promoter. Protein aggregation is scored by quantification of the EGFP fluorescent puncta in yeast. **(B-C)** Identification of killifish candidate proteins that do not aggregate (B) and candidate proteins that do aggregate (C) following galactose-inducible overexpression in yeast. White arrows: fluorescent puncta (aggregates). Scale bar: 5µm. **(D)** ‘Two-color’ experimental design to assess prion-like aggregate propagation in yeast. This design tests whether a brief overexpression (‘induction’) of a killifish candidate (C-terminally tagged with mRuby3 and under the control of a galactose-inducible promoter) can induce a cross-generational prion-like propagation of a constitutively expressed candidate (C-terminally tagged with eGFP and under the control of the GPD-promoter with single-copy number plasmid). The cross-generational effect was tested during ‘outgrowth’ (i.e., 200-fold dilution), highlighting aggregates that can stably propagate for 7-8 generations of yeast cells. Protein aggregation is scored by quantification of mRuby (red) or EGFP (green) fluorescent puncta in yeast. **(E)** Using the two-color system, killifish DDX5-EGFP aggregates were visible in both induction and outgrowth phases, indicating prion-like propagation. Representative of 4 independent experiments (quantified in F). White arrows: fluorescent puncta (aggregates). Scale bar: 5µm. **(F)** Quantification of DDX5 aggregates and prion-like seeding potential by calculating the fraction of EGFP positive cells with EGFP puncta during pre-induction, induction, and outgrowth. Mean values from 4-6 independent experiments are indicated by the black bar (total of 6 independent pre-induction measurements and four independent experiments with matched pre-induction, induction, and outgrowth measurements). Each dot represents the average value within each experiment (an average of 135 GFP-positive yeast cells were quantified for each strain under each experimental condition). Difference across groups were assessed using Student’s t-test. **(G)** The prion-like domain of DDX5 is necessary for propagation. Following overexpression of DDX5FL-mRuby3, DDX5ΔPrD-GFP forms aggregates (white arrows) in the induction phase but becomes diffuse (nuclear) in the outgrowth phase. Representative of 4 independent experiments (quantified in H). Scale bar: 5µm. **(H)** Quantification of DDX5 ΔPrD aggregation and prion-like seeding potential by calculating the fraction of EGFP positive cells with EGFP puncta during pre-induction, induction with DDX5 FL-mRuby3, and outgrowth. The black bar indicates mean values from 4-5 independent experiments (total of 5 independent pre-induction measurements and 4 independent experiments with matched pre-induction, induction, and outgrowth measurements). Each dot represents the average value within each experiment (an average of 135 GFP-positive yeast cells were quantified for each strain under each experimental condition). Difference across groups were assessed using Student’s t-test.

To test whether DDX5 aggregates are ‘infectious’, we used a 2-color system in which yeast cells are switched from raffinose to galactose medium to induce mRuby3-tagged proteins (induction), then switched back to raffinose medium and allowed to propagate for twenty generations (outgrowth). This allows preexisting aggregates to be diluted beyond detection levels and allows detection of protein aggregation seeded across generations onto the low expressed EGFP-tagged version (Figure 3D). We verified that killifish DDX5 protein inducibly aggregated in this two-color system (Figure 3E, induction, red puncta), whereas other control proteins did not (Figures S4A and S4B). Interestingly, killifish DDX5 tagged with EGFP (DDX5-EGFP) still formed puncta after propagation over multiple generations (Figure 3E, outgrowth, green puncta, and Figure 3F, quantification), consistent with an infectious prion-like behavior.

We asked whether the disordered prion-like domain within DDX5 is required for its aggregation and prion-like propagation. Truncating the entire IDR abolished DDX5 aggregation (and consequently any ability to seed) in this assay (Figure S4D). Interestingly, truncation of just the PrD domain still resulted in DDX5 aggregation (Figure 3G, induction, red puncta), similar to our observations in 293T cells.

However, this PrD truncation construct could not act as a template for prion-like seeding (Figures 3G, outgrowth, absence of green puncta, and Figure 3H, quantification, Figures S4C and S4D). Combined with our observations in 293T cells, these results identify separable molecular drivers of DDX5 aggregation and prion-like seeding.

### DDX5 aggregation exhibits additional characteristics of prion-like propagation

Does DDX5 exhibit additional hallmarks of prion-like propagation? One strong characteristic of prion-like propagation is its specificity. In our two-color seeding assay, DDX5 puncta were efficiently propagated when killifish DDX5-EGFP was expressed with DDX5-mRuby3 (see Figure 3). However, such an effect did not occur with the expression of mRuby3 protein or polyQ-mRuby3 aggregates, demonstrating that propagation is specific to DDX5 (Figures 4A, outgrowth, absence of green puncta, and Figure 4B, quantification, Figure S4D).

**Figure 4:**
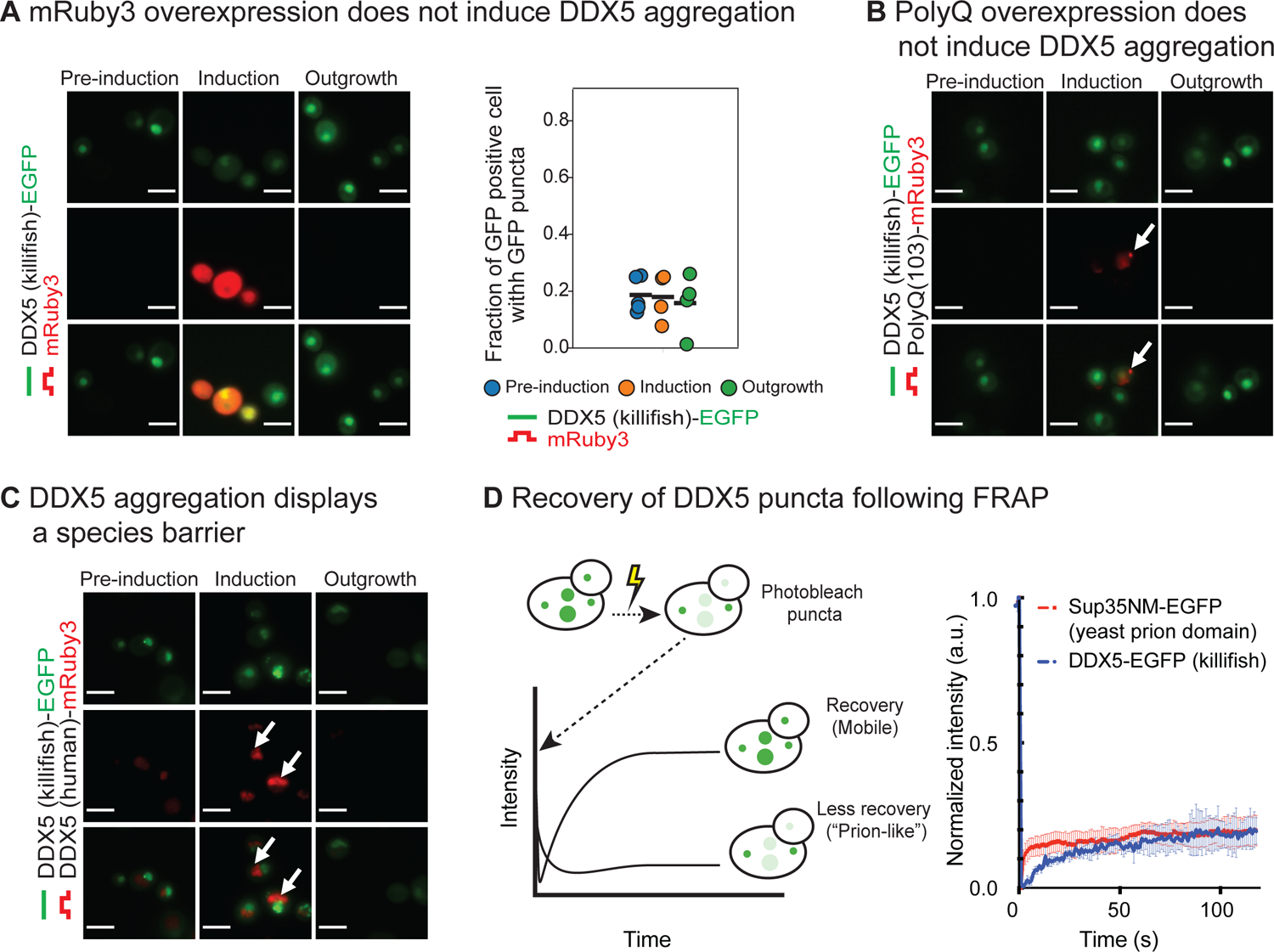
DDX5 exhibits additional prion-like characteristics. **(A)** Left: following expression of mRuby3 alone, killifish DDX5-EGFP remains diffuse (and nuclear-localized). Representative images of 4 independent experiments (quantified on the right). Scale bar = 5 µm. Right: Quantification of DDX5 aggregation and prion-like seeding potential by calculating the fraction of EGFP positive cells with EGFP puncta during pre-induction, induction, and outgrowth. Mean values from 4 independent experiments are indicated by the black bar. Each dot represents the average value within each experiment (an average of 135 GFP-positive yeast cells were quantified for each strain under each experimental condition). Difference across groups were assessed using Student’s t-test. **(B)** Following expression of the toxic and aggregation-prone PolyQ(103) repeat, killifish DDX5-EGFP remains diffuse (and nuclear-localized). White arrows: PolyQ(103)-mRuby3 aggregates. Mean values from 3 independent experiments are indicated by the black bar. Each dot represents the average value within each experiment (an average of 135 GFP-positive yeast cells were quantified for each strain under each experimental condition). Difference across groups were assessed using Student’s t-test. Scale bar: 5 µm. **(C)** DDX5 aggregation displays species barrier. Following overexpression of human DDX5-mRuby3, killifish DDX5-GFP remains diffuse (and nuclear-localized), highlighting species barrier – another characteristic of prion-like propagation. White arrows: DDX5 (human)-mRuby3 aggregates. Representative of 3 independent experiments. Scale bar: 5 µm. **(D)** Fluorescence recovery after photobleaching (FRAP) experiment to compare the recovery of killifish DDX5 (blue line) and yeast Sup35NM (a bona fide yeast prion domain, red line) puncta in yeast. Left panel: Prion-like proteins exhibit little fluorescence recovery by FRAP, indicating their aggregate nature. Right panel: Representative of 2 independent experiments. The trace represents mean +/- SEM of the normalized intensity of the photobleached area at each time point (10s interval with photobleaching performed at the 4^th^ frame).

Another key hallmark of prion-like aggregation, in contrast to other forms of protein misfolding, is a robust species barrier. Often, polymerization of a prion protein from one species does not efficiently promote the conversion of the same protein from another species, even though both can propagate independently (Santoso et al., 2000). Although killifish or human DDX5 could each aggregate efficiently, aggregation of human DDX5-mRuby did not spark prion-like propagation of killifish DDX5-GFP (Figure 4C). These data suggest a species barrier for DDX5 propagation.

A third characteristic of many prions is their ability to form relatively immobile aggregates. To investigate the dynamics of DDX5 aggregation, we used fluorescence recovery after photobleaching (FRAP) (Franzmann et al., 2018; Frederick et al., 2014; Maharana et al., 2018; Wu et al., 2006). In FRAP, traditional prion aggregates have poor recovery, consistent with their immobile nature (Figure 4D). We photobleached fluorescent puncta formed by overexpression of killifish DDX5-EGFP in live yeast cells. Fluorescence recovery was extremely slow and similar to the amyloid aggregates formed by the archetypical yeast prion protein Sup35 (Figure 4D). Collectively, these data suggest that DDX5 aggregates have several prion-like properties.

### DDX5 has phase separation properties that depend upon its disordered and prion-like domains

Some proteins with prion-like properties can also exhibit phase-separation behaviors, in which recombinant protein spontaneously forms large condensates within an aqueous solution (Boeynaems et al., 2018; Boeynaems et al., 2017; Franzmann et al., 2018; Hardenberg et al., 2020; Maharana et al., 2018; Molliex et al., 2015; Riback and Brangwynne, 2020; Riback et al., 2020; Sanders et al., 2020; van Mierlo et al., 2021). We characterized the phase separation properties of DDX5 *in vitro* by generating recombinant killifish DDX5 from *E. coli* (Figure S5A). Purified killifish DDX5 could easily phase-separate to form condensates under physiological protein and salt concentrations (1-10 µM and ∼150 mM, respectively) (Figure 5A). The formation of condensates was not due to molecular crowders or RNA (see STAR methods). These condensates were round, increased in size with time (Figure 5B), and they could be dissolved when elevating salt concentrations (Figure 5A). Moreover, condensates formed equally well with unlabeled protein (Figure 5B, left), establishing that the behavior was not an artifact of the fluorophore. Thus, killifish DDX5 exhibits a phase separation behavior *in vitro*, with the formation of large condensates.

**Figure 5:**
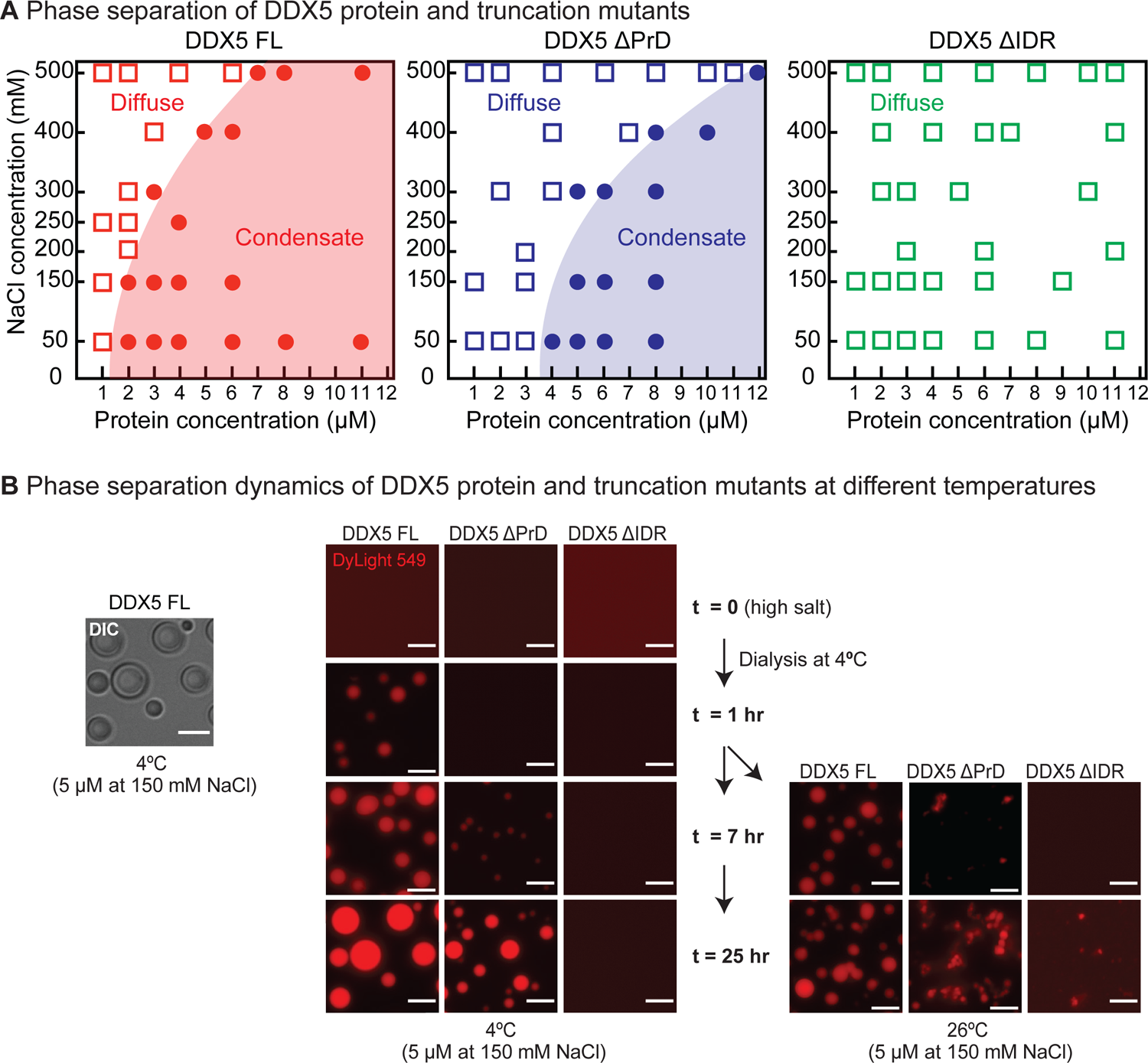
DDX5 phase separates and the formation of DDX5 condensates is dependent on its prion-like and disordered domains. **(A)** DDX5 can phase separate as a function of protein and NaCl concentration, and its prion-like domain (PrD) and intrinsically disordered region (IDR) are important to drive this behavior. Phase separation diagrams for purified killifish DDX5 recombinant protein full length (FL) and truncation mutants (ΔPrD and ΔIDR) under variable NaCl (y-axis, NaCl in mM) and protein concentrations (x-axis, in μM, range of protein concentration was estimated from mass spectrometry data). Empty squares: Diffuse state; full circle: Condensate state. **(B)** Left: a DIC representative image of DDX condensates. Right: temporal dynamics of the phase separation of DDX5 protein at various temperatures for DDX5 full length (FL) or different truncation mutants (ΔPrD and ΔIDR). Protein and salt concentrations are indicated. Red: DyLight 549 labeled C-terminally SNAP-tagged recombinant protein. Representative of 2 independent experiments. Experiments in (B) were performed with 5 μM protein at 150 mM NaCl. Scale bar: 5 µm unless indicated.

We then asked which domains of DDX5 are necessary for its *in vitro* phase separation properties. Truncation of the entire IDR completely abolished phase transition (Figure 5A). Truncation of only the PrD domain did not entirely abolish DDX5’s phase separation behavior, but it did shift the phase boundary (Figure 5A). Indeed, DDX5 ΔPrD mutant protein remained mostly diffuse under physiological concentrations (Figure 5A) and formed only small droplets in non-physiological (high protein concentration) conditions (Figure 5B). Thus, the PrD domain appears to be critical for phase separation at physiological conditions and for the formation of large condensates. Furthermore, given that the PrD domain is also essential for the propagation of aggregates *in vivo* in yeast (see Figure 3), our data are compatible with the possibility that the phase transition properties of DDX5 play an important role in driving its prion-like propagation.

### DDX5 condensates enhance activity but can mature into inactive, potentially infectious aggregates

DDX5 is an ATP-dependent RNA helicase. We measured DDX5 ATPase activity in the diffuse vs. condensate states. To this end, we isolated diffuse DDX5 that was still in pre-steady state condition (Figure 6A) by quickly dialyzing the protein in low salt buffer and removing any nucleated condensate that might have formed through centrifugation. We then performed an RNA-dependent ATPase activity assay on half of the diffuse DDX5 sample. In parallel, we waited for the other half DDX5 sample to mature, forming a mixture of diffuse proteins and condensates, and we performed the same RNA-dependent ATPase assay on this fraction (Figures 6A and Figure S6A). Interestingly, in conditions that favor condensate formation, DDX5 had a greater specific activity of ATP hydrolysis than when it was diffuse (Figure 6B). We next examined the ATPase activity of the DDX5 ΔPrD mutant protein. DDX5 ΔPrD did not form condensates at the concentrations used and did not exhibit a change in activity as a function of increased protein concentration (Figure S6B). The ATPase activity of DDX5 and DDX5 ΔPrD truncation mutant in the diffuse state were similar at saturating RNA concentrations (Figures S6C and S6D). These data suggest that the condensate phase may be physiologically important for DDX5 function.

**Figure 6:**
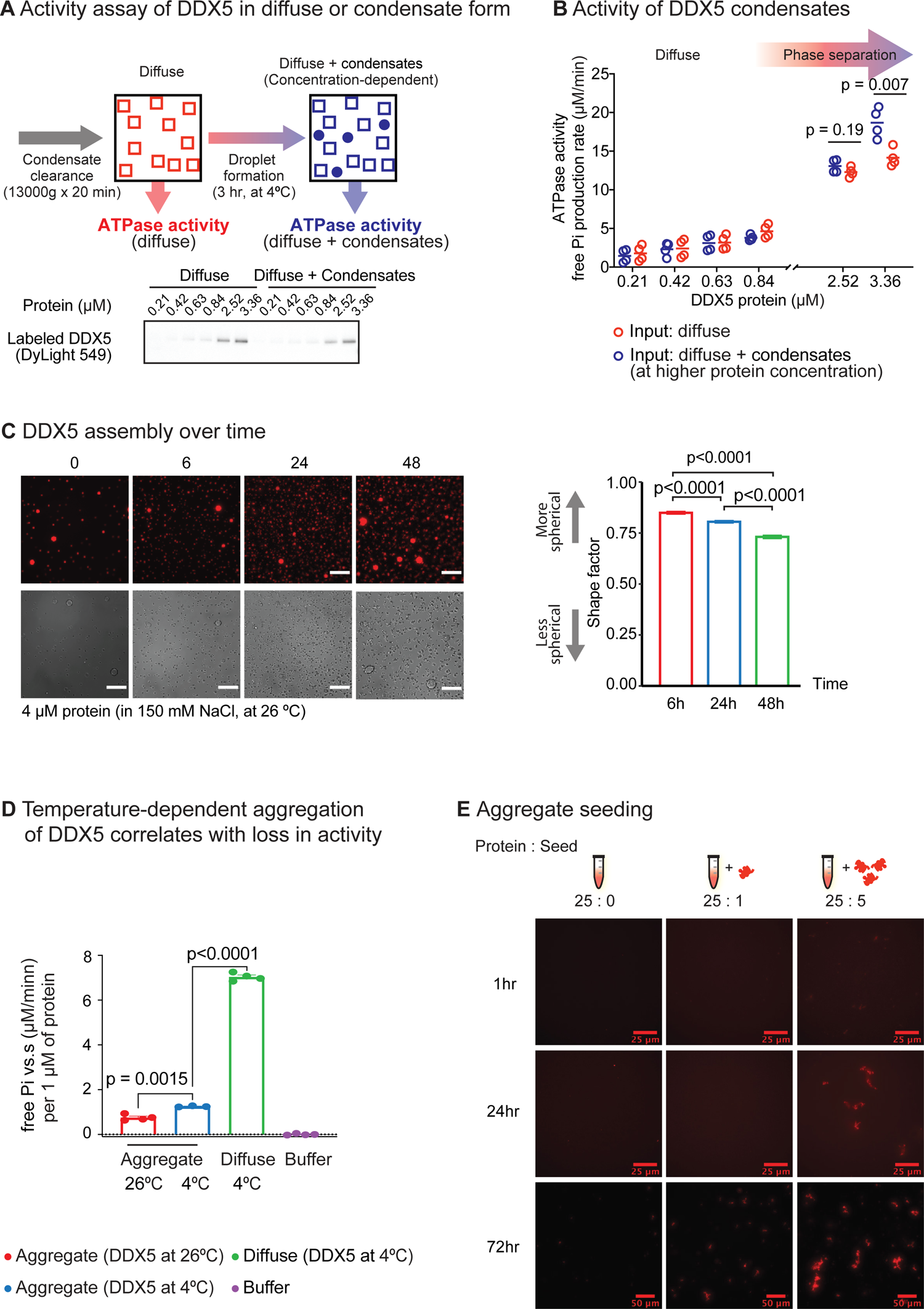
Characterization of the ATPase activity of DDX5 condensates and aggregates. **(A)** Experimental scheme used to measure the ATPase activity of the DDX5 in diffuse or condensates forms. Empty squares: diffuse state; full circle: condensates. Bottom panel: western blot to verify DDX5 protein concentration. **(B)** Assessment of ATPase activity for the killifish DDX5 protein in diffuse (red circles) or condensate (blue circles, above 2 μM) forms. ATPase activity is measured as a function of free phosphate (Pi) release rate (μM/min) at the indicated DDX5 protein concentrations (µM). Values of 4 replicates from one representative experiment are shown (corresponding to the western blot in (A)). Two independent experiments were performed and the result of the second is in Figure S6A. **(C)** Formation of less spherical (more disorganized) DDX5 aggregates over time. SNAP-tagged recombinant DDX5 was labeled with the DyLight 549 dye to allow visualization in the red channel (left panels). Quantification of the shape factor (corresponding to the sphericality of the condensate/aggregates) (right panel). P-values from a Student’s t-test. **(D)** ATPase activity experiment for the killifish DDX5 protein, comparing the activities of the aggregate (red or blue bars), diffuse forms (green bar), and buffer only (purple bar). ATPase activity is measured as a function of free Pi production rate (μM/min) per μM of protein. Representative of 2 many independent experiments (mean +/- SD from one experiment was shown). **(E)** Potential of DDX5 to form aggregates upon seeding with aggregated DDX5 protein (seed) at the indicated molar ratios (top) and time post-seeding (left). SNAP-tagged recombinant DDX5 was labeled with DyLight 549 dye to allow visualization in the red channel. The scale bar is either 25µm (top) or 50µm (bottom). Representative of 2 independent experiments.

Finally, we asked whether the physiological phase-separation properties of DDX5 could come at a price over time and potentially underlie the emergence of a prion-like seeding behavior. Over time DDX5 condensates matured into less spherical (more disorganized) aggregates (Figure 6C). Interestingly, while DDX5 condensates exhibited more ATPase activity than the diffuse form (see Figure 6B), these solid DDX5 aggregates had much less ATPase activity than the diffuse form (Figure 6D). These solid aggregates also efficiently seeded the formation of additional aggregates *in vitro* (Figure 6E). Collectively, these data suggest that the condensate form of DDX5 may enhance its physiological activity but could over time lead to irreversible aggregation, loss of function, and potentially seeding (Figure 7).

**Figure 7:**
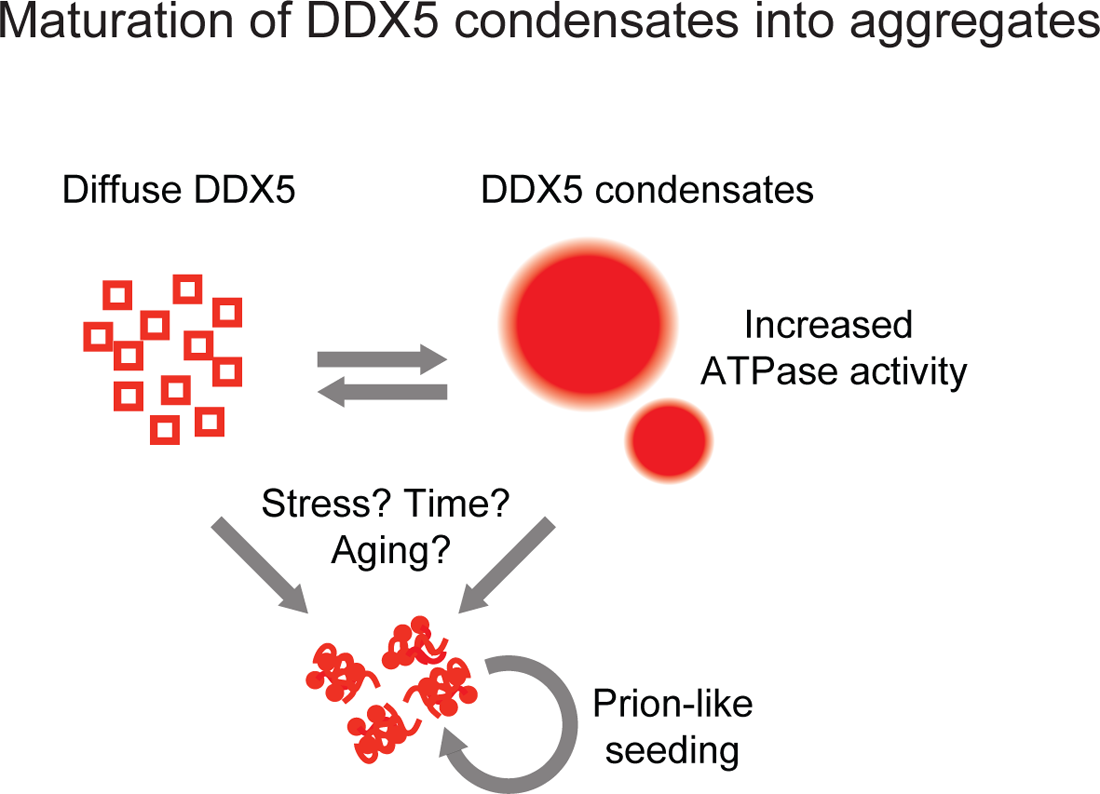
Putative model for the maturation of DDX5 condensate and the associated changes in activity Putative model depicting the maturation of DDX5 condensates (active) into aggregates (inactive) with prion-like seeding potential over time or during aging.

## DISCUSSION

Our study shows for the first time that many proteins aggregate increase in the vertebrate brain during normal aging. Protein aggregates were known to increase with age in *C. elegans* (Ben-Zvi et al., 2009; David et al., 2010; Huang et al., 2019; Labbadia and Morimoto, 2015; Lechler et al., 2017; Walther et al., 2015), but whether this applied to vertebrates which have more complex organs and tissues was not characterized quantitatively. Furthermore, while older killifish have been shown to exhibit aggregation of α-synuclein (Matsui et al., 2019) or ribosomal subunits (Kelmer Sacramento et al., 2020), our study reveals that many proteins that were not previously associated with diseases also aggregate during physiological aging. We showed that one of these proteins, the RNA helicase DDX5, can form aggregates in old brains both in killifish and in mice, indicating that protein aggregation is not only a consequence of the short killifish lifespan and that the aggregation of specific proteins is conserved throughout evolution. Proteins that aggregate during aging in the brain exhibit enrichment for disordered regions and prion-like (PrD) domains, suggesting a protein-autonomous mode of aggregate formation. However, the protein homeostasis environment of the brain during aging could also contribute to protein aggregation (Draceni and Pechmann, 2019; Gallotta et al., 2020; Muller et al., 2020) (see Chen, Harel, et al.).

DDX5 is an ATP-dependent RNA helicase that is involved in RNA splicing. DDX5 was previously found to associate with RNA foci in mammalian cells (Laurent et al., 2012). DDX5 naturally forms nuclear speckles, which are likely important for its normal function as a splicing helicase (Liao and Regev, 2021). Indeed, we show that DDX5 can self-assemble into droplets *in vitro*, and that these droplets have increased ATP hydrolysis activity, suggesting a physiological role for DDX5 nuclear speckles. However, this physiological property could also render DDX5 more susceptible to aggregation during aging. Abnormal DDX5 aggregation could result in RNA splicing defects and altered cellular functions. As DDX5 is involved in the splicing of Tau in Alzheimer’s disease (Kar et al., 2011), aggregation properties might contribute to defects during aging and underlie the age-dependency of neurodegenerative diseases.

Strikingly, DDX5 exhibits a previously unreported phase transition *in vitro* and prion-like behavior *in vivo* in yeast. Other RNA-binding proteins (from yeast and mammals) have been recently shown to exhibit phase separation behavior (FUS, TDP-43, SUP35, etc.), and their PrD domains are important for this transition (Franzmann et al., 2018; Guo et al., 2018; Maharana et al., 2018; Tauber et al., 2020; Wang et al., 2018). We find that the PrD domain of DDX5 is critical for phase transition and prion-like properties. These prion-like phase separation properties could contribute to the spread of cognitive defects during aging and neurodegenerative diseases.

## ACKNOWLEDGMENTS

We thank Steven Boeynaems, Serena Sanulli, and Reut Shalgi for critical reading of the manuscript. We thank the Brunet, Jarosz, and Harel labs, and in particular Olivia Zhou, for stimulating discussion and feedback on the manuscript. We thank T. Kelly Rainbolt and Judith Frydman for their guidance on protein homeostasis. We thank Christine Yeh for their help with formulation of the initial mass spectrometry analyses. We thank Stanford’s mass spectrometry facility, in particular Christopher M. Adams and Ryan Leib, for processing the samples. We thank Parag Mallick and Josh Elias for feedback on mass spectrometry analyses. We thank Kiran Chandrasekher and Ray Futia for independent and blinded quantification of protein aggregation from yeast microscopy images. We thank Anupam Chakravarty for helpful discussions on DDX5 purification and *in vitro* experiments. We thank Susan Murphy (Stanford) and Ashayma Abu-tair (HUJI) for help with killifish maintenance. This work is supported by NIH RF1AG057334 (A.B., D.F.J), R01AG063418 (A.B., D.F.J.), the Stanford Alzheimer’s Disease Research Center and the Zaffaroni Alzheimer’s Disease Translational Program (A.B., D.F.J.), the Stanford Brain Rejuvenation Project (A.B.), the Glenn Foundation for Medical Research (A.B.), the Stanford Systems Biology Seed Grant (I.H., Y.R.C., and I.Z.), Abisch-Frenkel Foundation 19/HU04 (I.H.), Zuckerman Program (I.H.), NIH 1R21AG063739 (I.H.), ISF 2178/19 (I.H.), Israel Ministry of Science 3-17631 and 3-16872 (I.H.), Moore Foundation GBMF9341 (I.H.), BSF-NSF 2020611 (I.H.), and Israel Ministry of Agriculture 12-16-0010 (I.H). I.H. was supported by a Damon Runyon, Rothschild, and Human Frontiers long-term post-doctoral fellowships. Y.R.C. was supported by a Stanford Graduate Student Fellowship.

## AUTHOR CONTRIBUTIONS

I.H., Y.R.C., I.Z., A.B., and D.F.J. designed the mass spectrometry experiments. I.H. isolated killifish organs, Y.R.C. and I.Z. isolated samples for mass spectrometry analysis with the help of I.H. and performed silver stain analysis. Y.R.C. designed and implemented mass spectrometry data analysis and performed domain enrichment analyses with input from I.H., P.P.S., A.B. and D.F.J.. I.H. generated mammalian constructs, performed and analyzed the immunocytochemistry experiments and 293T aggregation experiments, generated the DDX5 CRISPR knock-out, and the DDX5 antibody. Y.R.C. generated yeast constructs with help from I.H. and B.E.M, and performed and analyzed the yeast aggregation and FRAP experiments. Y.R.C. generated and purified recombinant killifish DDX5 proteins and performed and analyzed phase separation and activity assays. P.P.S. conducted PCA and GO term analyses. P.N.N. performed and analyzed DDX5 immunocytochemistry experiments in mice. U.G. and G.A. performed and analyzed immunocytochemistry experiments in killifish and western-blot for DDX5, respectively, under the supervision of I.H. I.H. generated the transgenic DDX5 construct, and W.W. generated the transgenic killifish overexpressing DDX5 under the supervision of A.S.A.. E.M. and A.M. provided young and old killifish brain sections. K.H. performed an initial analysis of the mass spectrometry data. S.Y. helped with sequence verification and cloning of killifish constructs. I.H., Y.R.C., D.F.J. and A.B. wrote the paper, and all the authors commented on the manuscript.

## STAR METHODS

### Key Resources Table

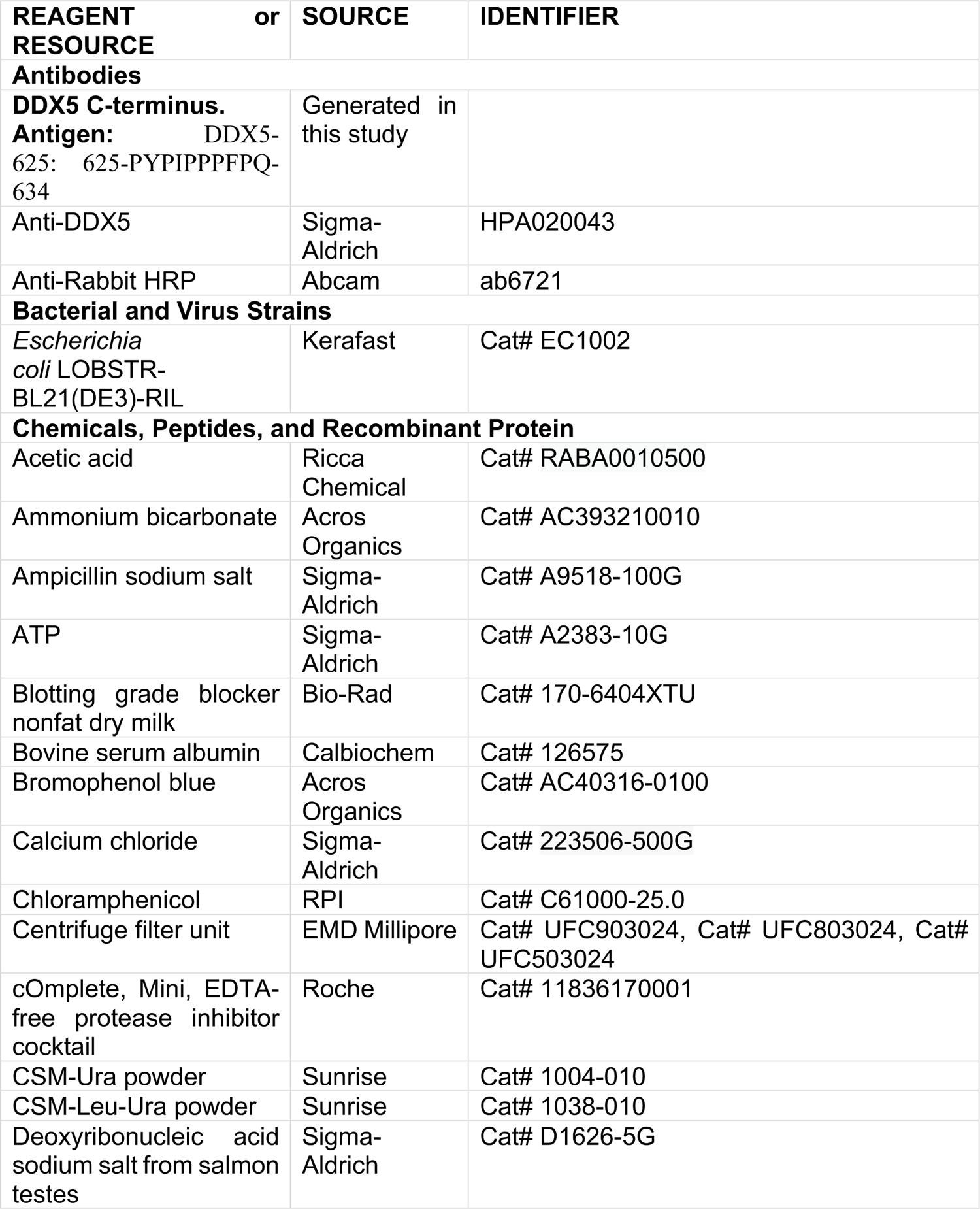
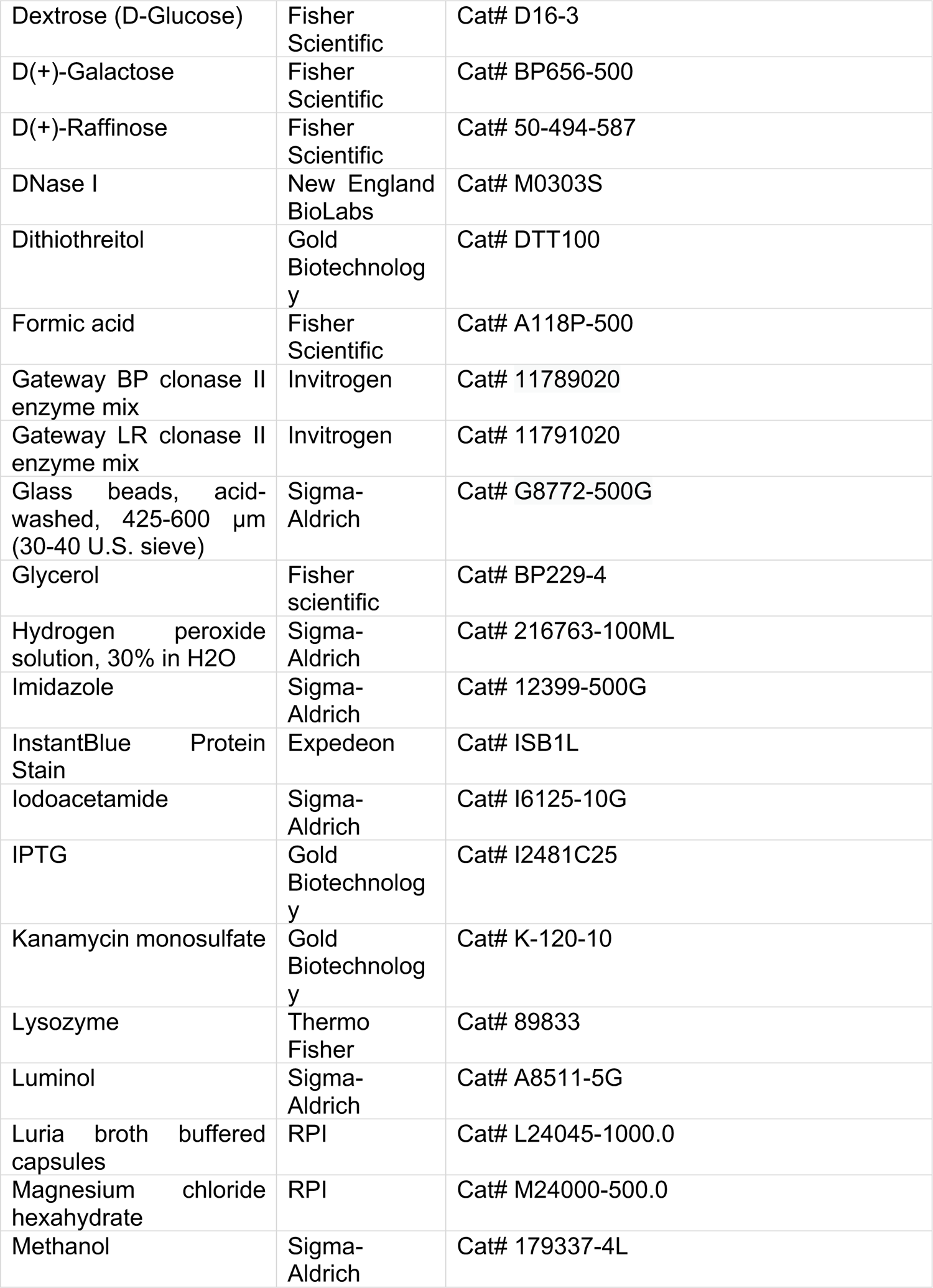
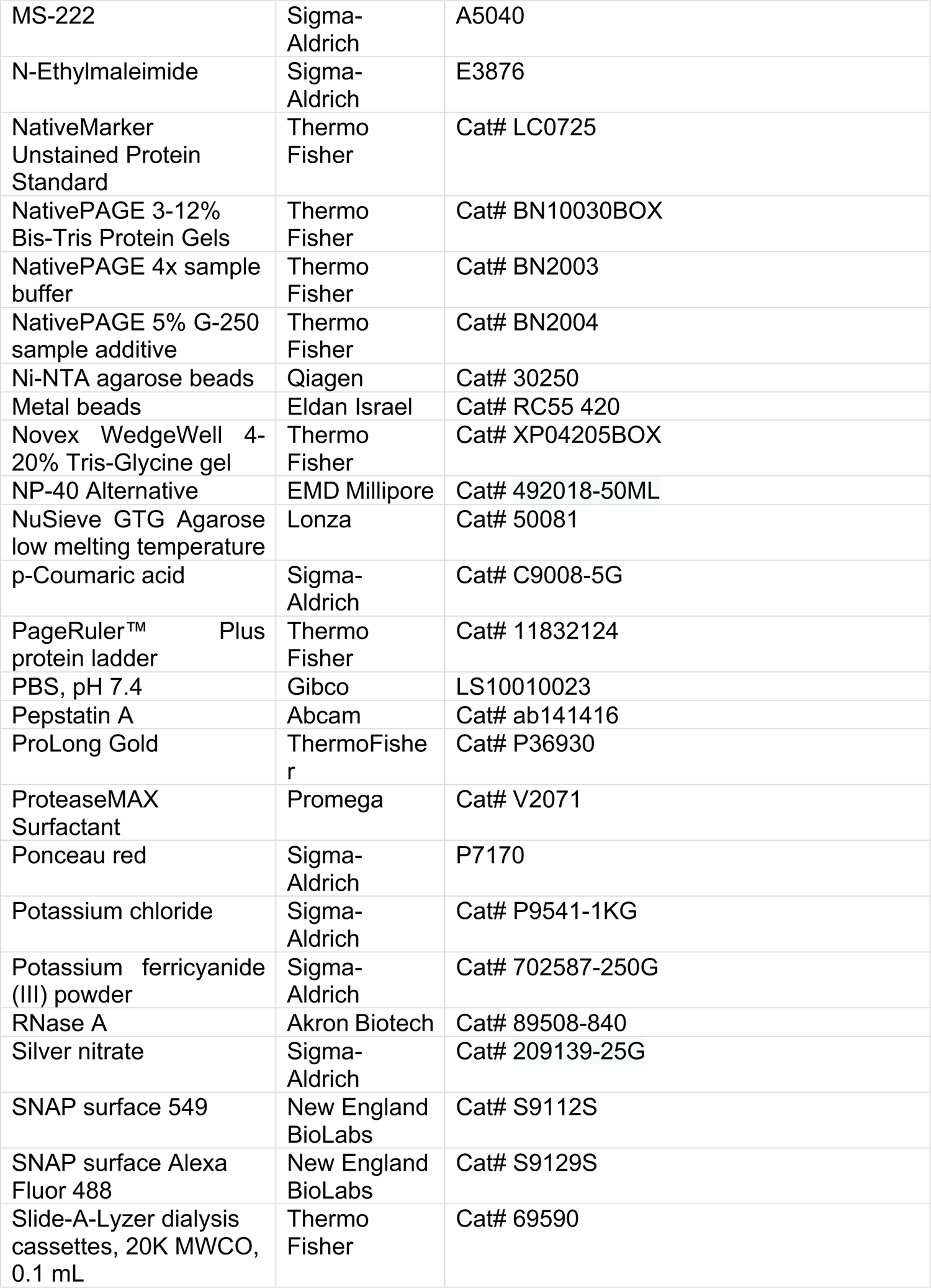
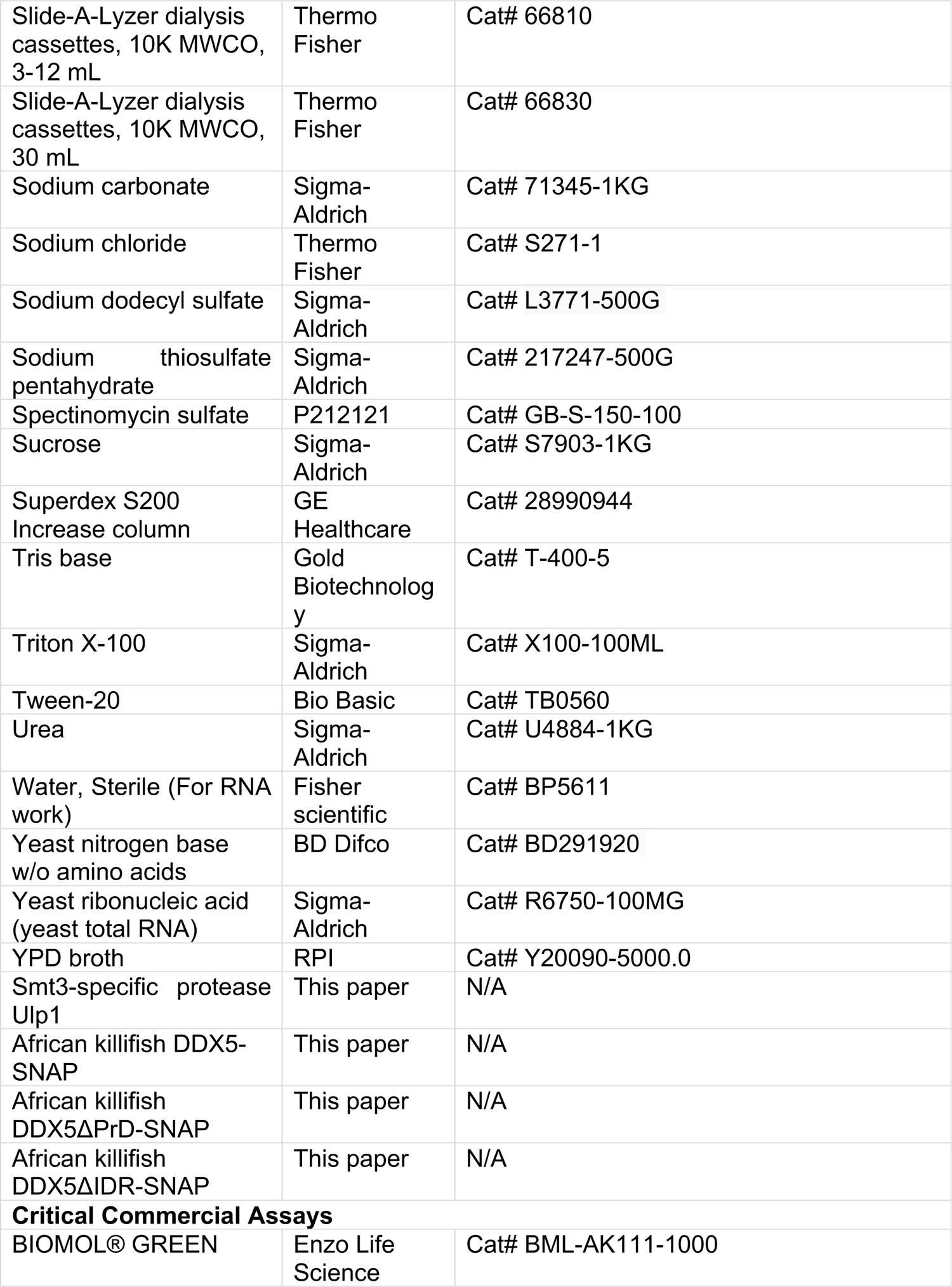
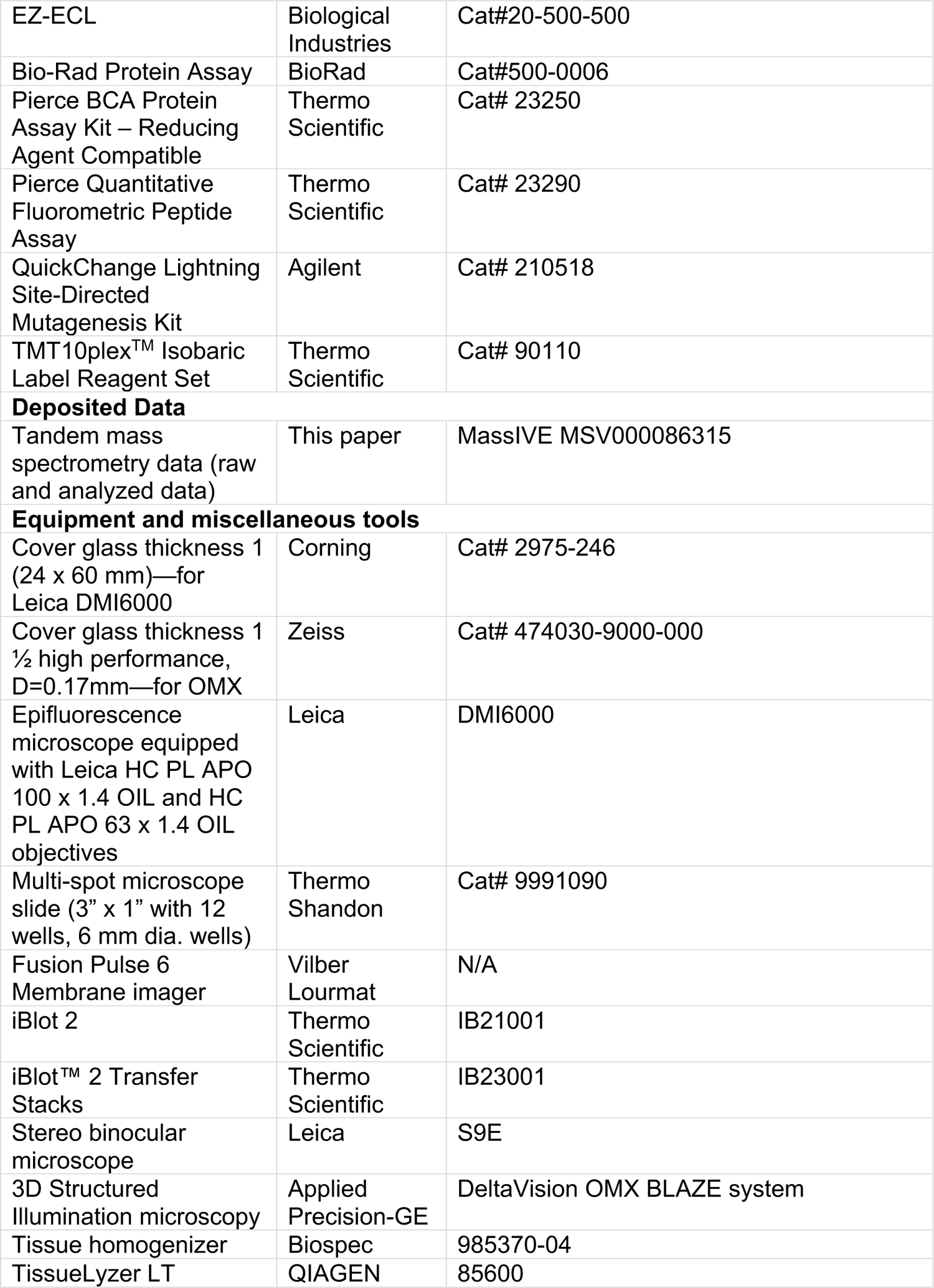
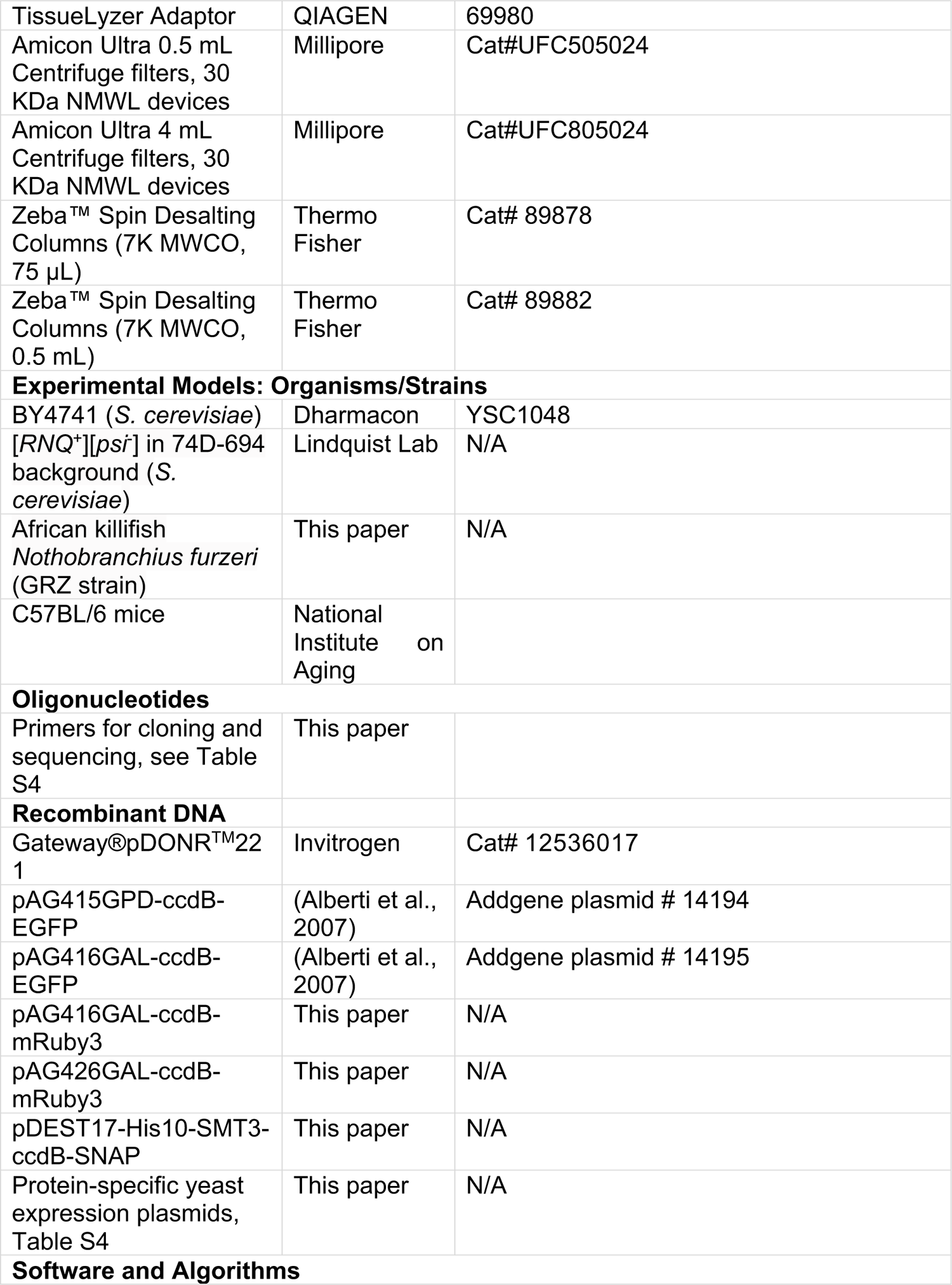
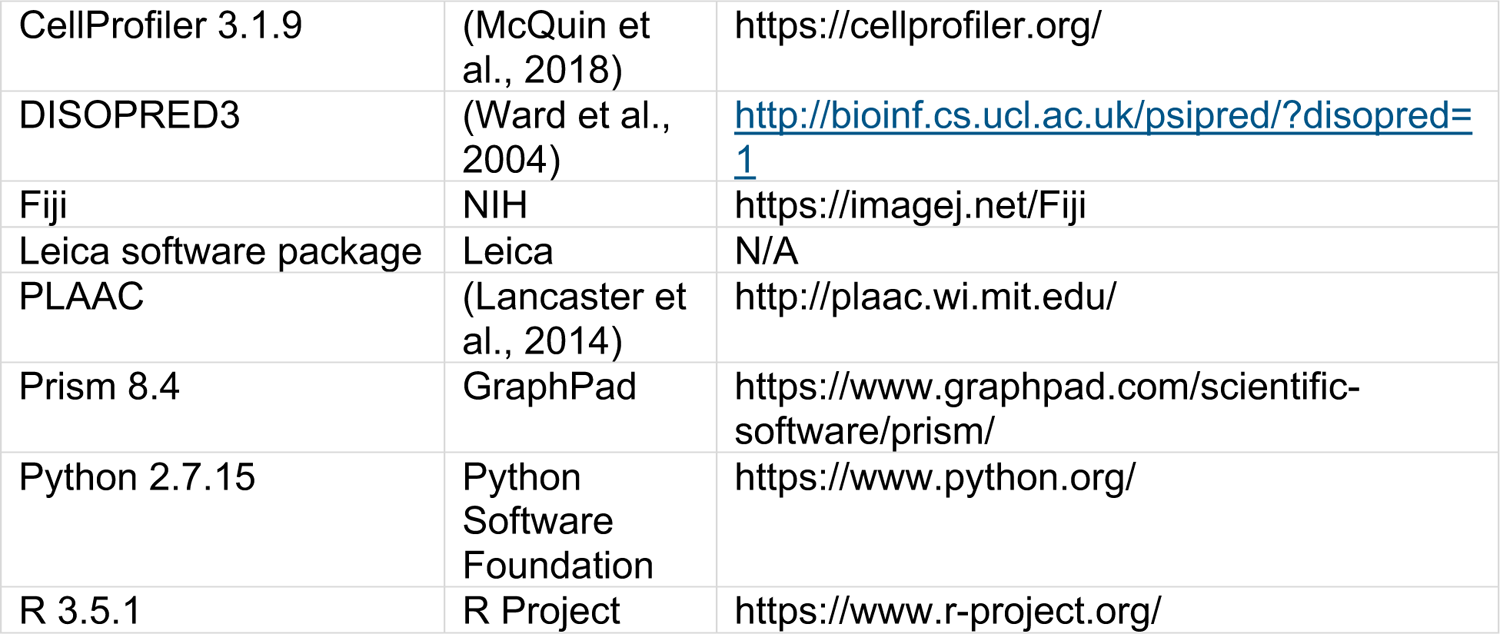

### Lead Contact for Reagent and Resource Sharing

Anne Brunet and Dan Jarosz

### Data and Code availability

All raw mass spectrometry reads as well as processed datasets can be found in the MassIVE database (https://massive.ucsd.edu/ProteoSAFe/static/massive.jsp) under ID MSV000086315. The codes and results supporting to the current study are available in the Github repository for this paper https://github.com/ywrchen/killifish-aging-aggregates.

### Experimental Model and Subject Details

#### S. cerevisiae

*S. cerevisiae* strains were obtained from the sources indicated (Key Resources Table). All *S. cerevisiae* strains were stored as glycerol stocks at −80°C. Before use, strains were either revived on YPD (10 g/L yeast extract, 20 g/L dextrose, 20 g/L peptone, sterilized by autoclaving) or on defined medium (2% glucose, 6.7 g/L yeast nitrogen base without amino acids, 20 mg/L histidine, 120 mg/L leucine, 60 mg/L lysine, 20 mg/L arginine, 20 mg/L tryptophan, 20 mg/L tyrosine, 40 mg/L threonine, 20 mg/L methionine, 50 mg/L phenylalanine, 20 mg/L uracil, 20 mg/L adenine, sterilized by autoclaving) as necessary. Antibiotics, or defined drop-out media were used as indicated to maintain plasmid selection. All strains were grown at 30°C unless otherwise indicated.

Yeast strains expressing exogenous killifish proteins were generated by transforming laboratory strain BY4741 (either fresh mid-exponential cells or frozen chemically competent cells) with yeast expression plasmids that encoded proteins of interest. Yeast transformation was carried out using a standard lithium-acetate protocol. First, cells were inoculated into 25 mL of liquid rich medium (YPD) and grown to saturation overnight on a shaker at 200 r.p.m and 30°C. The cells were then diluted by 25-fold into 500 mL of liquid rich media (YPD) and regrown on a shaker at 200 r.p.m and 30°C. Once the culture reached mid-exponential phase (OD_600_ ∼0.4 - 0.6), the cells were harvested by centrifugation at 2000xg for 5 min and washed twice in an equal volume of sterile water. The cells were either used directly for yeast transformation or further processed to generate competent cells. To generate chemically competent cells, cell pellets were resuspended in 5 mL of filtered sterile frozen competent cell solution (5% v/v glycerol, 10% v/v DMSO), and 50 µL aliquots were generated in 1.5 mL microcentrifuge tube and stored at −80°C. To ensure good survival rates, aliquots were slowly frozen either using Mr. Frosty freezing container (Thermo Scientific Cat# 5100-0001) or Styrofoam box padded with Styrofoam chips or newspaper (to reduce air space around sample). For yeast transformation, competent cells were thawed in 37°C water bath for 15-30s then centrifuged at 13000xg for 2min to remove supernatant and resuspended in a transformation master mix (260 μL PEG 3500 50% (w/v), 36 μL 1 M Lithium acetate, 50 μL denatured salmon sperm carrier DNA (2 mg/mL), 14 μL plasmid DNA (0.1-1 μg total plasmid), and sterile water to a final volume of 360 μL). Cells were incubated in the transformation master mix at 42°C for 45 min. Following incubation, cells were harvested, resuspended in 1 mL sterile water, and ∼100 μL was plated on selective medium and grew at 30°C. Successful transformants typically appeared in 2-3 days and were further propagated in defined liquid drop-out medium (omitted nutrient depending on the plasmid being selected) and stored as glycerol stocks in −80°C.

#### Mice

All mice used in this study were male C57BL/6 mice obtained from the NIA Aged Rodent colony. Mice were habituated for more than one week at Stanford before use. At Stanford, all mice were housed in the Comparative Medicine Pavilion or the Research Animal Facility II, and their care was monitored by the Veterinary Service Center at Stanford University under IACUC protocol 8661.

#### African turquoise killifish strain, husbandry, and maintenance

The African turquoise killifish (GRZ strain) were housed as previously described (Harel et al., 2015a). Fish were housed at 26°C in a central filtration recirculating system with a 12 hr light/dark cycle at the Stanford University facility (Aquaneering, San Diego) or at the Hebrew University of Jerusalem (Aquazone ltd, Israel). In both facilities, fish were fed twice a day on weekdays and once a day on weekends with Otohime Fish Diet (Reed Mariculture). In these conditions, killifish lifespan was approximately 6-8 months. The *TERT^Δ8/Δ8^* loss-of-function allele (Harel et al., 2015a) was maintained as heterozygous (due to fertility issues in homozygous) and propagated by crossing with wild-type fish. Following genotyping at the age of one month (Harel et al., 2015a), fish were single-housed. All turquoise killifish care and uses were approved by the Subcommittee on Research Animal Care at Stanford University (IACUC protocol #13645) and at the Hebrew University of Jerusalem (IACUC protocol #NS-18-15397-2).

## Method Details

### Isolation of Tissue Lysate (TL) and Aggregate-Enriched (AGG) fractions from brain tissues

This protocol is described in more depth in Chen, Harel et al for all tissues. Briefly, whole brains of 3 young (3.5 months) wild-type, 3 old (7 months) wild-type, and 3 old (7 months) *TERT^Δ8/Δ8^* mutant (Harel et al., 2015a) male fish were collected and snap-frozen in liquid nitrogen. All procedures were conducted at 4°C unless stated otherwise. Brains were homogenized using a tissue homogenizer in 100 µL of buffer A (30 mM Tris-Cl pH = 7.5, 1 mM DTT, 40 mM NaCl, 3 mM CaCl_2_, 3 mM MgCl_2_, 5% glycerol, 1% Triton X-100, protease inhibitor cocktail tablet used at 1x the manufacturer recommended concentration (Roche cOmplete^TM^ EDTA-free Protease Inhibitor Cocktail, Cat# 11697498001). Homogenization was performed in round-bottom tube (2 mL corning cryogenic vials) to ease lysis and the resulting sample was transferred to 1.5 mL Eppendorf tube for the first centrifugation spin. Lysate was spun at 800 g for 10 min (spin 1) to remove cell debris (Eppendorf Centrifuges 5430 with Eppendorf FA-45-48-11 30-spot 45-degree fixed angel rotor). Supernatants were transferred to a new Eppendorf tube and treated with 100 µg/mL RNase A (Akron Biotech, 89508-840), and 100 µg/mL DNase I (New England Bio, Cat# M0303S) for 30 min on ice. Samples were spun at 10,000 g for 15 min (spin 2 in the same Eppendorf FA-45-48-11 rotor) and the resulting supernatant is the tissue lysate (TL) fraction. A 25 µL aliquot of the TL was kept in a separate tube for protein quantification and mass spectrometry analysis. For isolation of the aggregate (AGG) fraction, all the remaining TL was loaded onto the top of a 1 mL 40% sucrose pad and additional ∼ 750 µL (adjusted to balance all ultra-centrifuge tubes) of buffer A was layered on the top in ultra-centrifugation tube (Beckman Coulter Ultra-Clear centrifuge tubes, thinwall, 2.2 mL, 11 x 34 mm, catalog # 347356). The samples were separated through ultracentrifugation for 1 hr at 200,000 g (spin 3 at 49,000 r.p.m. in Beckman TLS-55 rotor). Top layers of supernatants were removed, leaving 15-20 µL of liquid at the bottom. An additional 30 µL of buffer A was added to rigorously re-suspend the pellets. Protein concentration for TL and AGG samples was assessed using standard BCA kit (Pierce BCA Protein Assay Kit – Reducing Agent Compatible, catalog #23250).

### Mass spectrometry sample preparation and analysis

Following tissue lysate (TL) extraction and aggregate (AGG) isolation, samples were re-suspended in 8M urea and ProteaseMAX (Promega) and were subsequently subjected to reduction (with 10 mM DTT for 30 min at 55°C) and alkylation (with 30 mM acrylamide for 30 min at room temperature) followed by trypsin digestion (1:50 concentration ratio of sequencing grade trypsin to total protein overnight at 37°C followed by quenching with 25 µL of 50% formic acid to pH below 3.0). The digested peptides from different samples of the same organ were separately quantified and equal amount of peptide samples were labeled with TMT10plex mass tag before mass spectrometry analysis. In particular, equal amount of TL or AGG (3-10 µg of peptide depending on the organ/tissue) from each sample was labeled with 9 different TMT-10plex tags accordingly to the manufacturer’s protocols (cat# 90110, Thermo Scientific). The same mass tag and sample assignment was maintained throughout the entire study (old brain samples were labeled with TMT^10^-126, TMT^10^-127N, and TMT^10^-127C respectively; young brain samples were labeled with TMT^10^-128N, TMT^10^-128C, and TMT^10^-129N respectively; and old *TERT^Δ8/Δ8^* brain samples were labeled with TMT^10^-129C, TMT^10^-130N, and TMT^10^-130C respectively). One ninth of each sample (equal amount of peptide across all TL or AGG samples from a tissue/organ) were pooled after trypsin digestion and labeled with the 10^th^ TMT tag to serve as the reference channel for internal normalization. Post-labeling, each set of samples were further cleaned up with C18 peptide desalting columns. Samples were subjected through high pH reverse phase fractionation into 3 or 4 fractions and all fractions were run independently on an Orbitrap Fusion (Thermo Scientific) mass spectrometer coupled to an Acquity M-Class nanoLC (Waters Corporation). Data searches were conducted with killifish proteome downloaded from NCBI (available in GitHub repository https://github.com/ywrchen/killifish-aging-aggregates and MassIVE dataset MSV000086315 for this paper). Mass spectra were analyzed using Proteome Discoverer v2.0 (Thermo Scientific) for MS3 quantification of tandem mass tag reporter ions and the Byonic v2.6.49 search algorithm node for peptide identification and protein inference. Briefly, a typical mass spectrometry analysis allowed for fully tryptic digestion with up to two missed cleavages. A 12 ppm mass accuracy was tolerated for precursor and MS3 HCD fragments, *i.e.*, reporter ions, and 0.3 Da mass accuracy for CID fragmentation at the MS2 level. Static modifications include cysteine carbamidomethylation and TMT labels on peptide N-termini and lysine residues. Oxidation of methionine and deamidation of aspartate and glutamine were considered as dynamic modifications. Peptides and proteins were cut at the 1% FDR level using the Byonic node. Reporter ion intensities were normalized against a pooled sample containing each of the other samples in a given sample run and reported relative to these pooled samples. These ratios were exported for analysis at both the protein and PSM (Peptide Spectrum Match) level. The mass spectrometry raw data, summary table of the reporter ion ratios at protein and PSM levels, as well as protein sequence FASTA files use for the search were deposited to MassIVE with a dataset id as MSV000086315.

### Annotation of the NCBI genes models of turquoise killifish

We used the African turquoise killifish (*N. furzeri*) NCBI annotation release 100 for our analysis. The majority of the gene models in this annotation only have a locus number. Therefore, we re-annotated all the killifish gene model names based on orthology analyses with 40 species including mammals, fish and invertebrates. We selected the consensus symbols for the locus assigned by NCBI as our final symbols and re-annotated the genes using a naming scheme Gene_Name(n of m) if there were duplicates in the killifish genome. For most of the analyses, human homologs were reported or used in database search, unless otherwise noted. The annotation and human homologs used in this killifish study can be found in Table S1C.

### Mass Spectrometry Data Normalization and Analysis of Age-associated Changes

The target protein results including the reporter ion ratios and total number of spectra assigned to peptides from this protein (# PSMs) were further processed to infer the abundance of each protein in each sample. First, the human contaminants were removed. Next, the protein abundance of each sample was inferred from its PSM contribution (or equivalently each TMT10plex tag), calculated by multiplying the total number of PSMs for a protein by the fraction of reporter ion signal that came from this channel (ratio of query channel divided by sum of the ratios across all channels, note that the TMT-131 was the normalization channel and contributed as 1 to the overall signal). Because equal amounts (by mass) of peptides were loaded in each channel, we normalized the sum of PSMs for all proteins in a channel to a constant of 100,000. The resulting normalized counts of PSMs for a protein in a sample represent the final reported protein abundance (PSMsNorm). We log2-transformed the protein abundance (log2_PSMsNorm) and the resulting protein abundance for each tissue approximately followed a normal distribution (Cite Chan, Harel). We then used the log2-transformed protein abundance to perform a parametric statistical test (Student’s t-test), as is often done for proteomics datasets (Klann et al., 2020; Li et al., 2019; Mirzaei et al., 2017; Navarrete-Perea et al., 2018; Nusinow et al., 2020; Zhang and Elias, 2017).

The age-associated changes in a protein in either tissue lysate or aggregate fraction were calculated as the fold change in the average abundance of a protein between the two age/disease groups. Both fold change (i.e. OvY_FC for the fold change of old divided by young) and log2-transformed fold change (i.e. OvY_logFC was the log2-transformed fold change of old divided by young) are reported in Table S1A. Age/disease group differences were assessed using a Student’s t-test (p-values from this test referred to as OvY_pval). The p-values were not corrected for multiple hypothesis testing, as is often done for proteomics datasets (Klann et al., 2020; Li et al., 2019; Mirzaei et al., 2017; Navarrete-Perea et al., 2018; Nusinow et al., 2020; Zhang and Elias, 2017).

We defined the term ‘aggregation propensity’ (PROP) to infer the intrinsic likelihood of a protein to aggregate, scored by dividing the abundance of a protein in the aggregate fraction (AGG) by its tissue lysate (TL) abundance. Note that this metric can only be reported when a protein is identified in both the TL and AGG fractions. Proteins that were only detected in AGG but not in TL make up about 0.8-1.7% (median 1.3%, average 1.2%) of total AGG signal and their abundance change was analyzed at the AGG level only. Because each channel represents the tissue sample from an individual fish, we reported the aggregation propensity of a protein for each sample. The age-associated changes in aggregation propensity (i.e. OvY_prop_FC) were calculated as the fold change in the average aggregation propensity of a protein between the two age/disease groups. The log2-transformed fold change (i.e. OvY_prop_logFC) was reported as well. Student’s t-tests were performed on the log2-transformed aggregation propensities of the different conditions (Young, Old, Old *TERT^Δ8/Δ8^*mutants). The resulting p-values were reported (OvY_prop_pval) to assess whether the changes between conditions were statistically significant.

### Principal component analysis (PCA)

Standardized log2-transformed normalized protein abundance (peptide spectra account for each protein) for young, old, and *TERT^Δ8/Δ8^* mutant brain samples (TL and AGG) were used as input for principal component analysis (PCA). PCA was performed in R (version 3.5.0) using autoplot function implemented in R package ggfortify (version 0.4.6) and plotted using ggplot2 (version 3.1.1).

### Gene Ontology Enrichment Analysis

Enriched Gene Ontology (GO) terms associated with proteins from old versus young brain samples (AGG and TL) were identified using Gene Set Enrichment Analysis (GSEA) implemented in R package clusterProfiler (version 3.10.1) (Yu et al., 2012). All the proteins identified in the brain were ranked and sorted in descending order based on multiplication of log_2_ transformed fold change and –log_10_(p-value). This metric prioritizes proteins with both large absolute fold changes and small p-values. Note that due to random seeding effect in GSEA, the exact p-value and rank of the enriched terms may differ for each run. These random seeds did not qualitatively affect the enrichment analyses.

### Statistics for enrichment for PLAAC and DISOPRED

We calculated the normalized prion score (NLLR) predicted by PLAAC (Lancaster et al., 2014) and the fraction of disordered residues with DISOPRED 3 (Jones and Cozzetto, 2015) for each protein that showed significant age-associated increase in aggregation in the brain and other tissues (paired t-test p-value <0.05 and t-value of log2 transformed fold change of AGG or PROP from old over young animals greater than 0.75). The p-values were not corrected for multiple hypothesis testing, as is often done for proteomics datasets (Klann et al., 2020; Li et al., 2019; Mirzaei et al., 2017; Navarrete-Perea et al., 2018; Nusinow et al., 2020; Zhang and Elias, 2017). We considered a protein to contain a prion-like domain (PrD) if it had a positive NLLR score in the primary sequence by PLAAC. We considered a protein to contain an intrinsically disordered region (IDR) if the fraction of disordered residues for that protein was greater than 0.3. Residues are considered disordered if their scores were greater than 0.5 based on DISOPRED 2 prediction. We then performed a Fisher’s exact test on the age-associated aggregates that contain either PrD or IDR, respectively, in comparison to the total detected aggregates in each tissue. The p-values were reported.

### CRISPR/Cas9 target prediction and gRNA synthesis

CRISPR/Cas9 genome-editing was performed according to (Harel et al., 2015b; Harel et al., 2016). In brief, for targeting DDX5, we identified conserved regions in the coding sequence using multiple vertebrate orthologs using http://genome.ucsc.edu/ (Figure S3). Conserved regions that were upstream of functional or active protein domains were selected for targeting in exon 5 and 7 of *DDX5*. gRNA target sites were identified using CHOPCHOP (https://chopchop.rc.fas.harvard.edu/) (Montague et al., 2014), and were as follows (PAM sites are in bold, in modified nucleotides, from C to G, are in square brackets): Exon5: [C]GACCATGTCTTTTCCACTC**AGG;** G[C]ATCGCTCAAACGGGGTCT**GGG;** GGGTGGCCCATTGTCCTGAG**TGG.** Exon7: GGTGGCTGCTGAGTACGGCA**GGG;** GGTGCATGTACTCTTGAGAC**GGG;** GGATGTACTCTTGAGACGGG**AGG;** GGTGGCACCAACCCGTGAGC**TGG.** Hybridized oligonucleotides, according to (Harel et al., 2015b; Harel et al., 2016), were used as an *in vitro* transcription template. gRNAs were *in vitro* transcribed and purified using the MAXIscript T7 kit (ThermoFisher # AM1312), according to the manufacturer’s protocol.

### Production of Cas9 mRNA

Experiments were performed according to (Harel et al., 2015b; Harel et al., 2016). The pCS2-nCas9n expression vector was used to produce Cas9 mRNA (Addgene, #47929) (Jao et al., 2013). Capped and polyadenylated Cas9 mRNA was in vitro transcribed and purified using the mMESSAGE mMACHINE SP6 ULTRA (ThermoFisher # AM1340).

### Microinjection of turquoise killifish embryos and sequencing of targeted sites

Microinjection of turquoise killifish embryos was performed according to (Harel et al., 2015b; Harel et al., 2016). Cas9-encoding mRNA (300ng/μL) and gRNA (30ng/μL) were mixed with phenol-red (2%) and co-injected into one-cell stage fish embryos. Three days after injection, genomic DNA was extracted from 5–10 pooled embryos. The genomic area encompassing the targeted site (∼600bp) was PCR-amplified, using the following primer sequences: Exon5F1: TTGGATAAGACACTCACTGCCA; Exon5F2: ACACTCACTGCCAAAGTCTTCC. Exon5R1: CTGATGTTGGATGTGAACGACT; Exon5R1: GTGAACTTACAATAGGCCCGTC. Exon7F1: CCAGAAAAGAGAGAGCGAGTTT; Exon7F2: TCATTATCGGACTCAGTTGCTT; Exon7R1: CTATCTTGCGGATCTGAGGTTC; Exon7R2: TCCGATACACGGTTTTTGAATA. DNA sequencing was used for analysis. Perturbing the DDX5 gene led to embryonic lethality in the F0 generation, resulting in failure to obtain germline-transmission and an F1 generation.

### Generation of DDX5-GFP transgenic fish

The *Tg(DDX5p:DDX5-GFP)* transgenic animals were generated at the Stowers Institute via the Tol2 system (Harel et al., 2016; Valenzano et al., 2011). A 7.8 kb cassette that includes 5.2 kb *DDX5* promoter sequence and 2.3 kb encoding a DDX5-GFP fusion protein was cloned into pDest-Tol2 vector (Kwan et al., 2007) through Gibson assembly (#E5510, NEB). Gibson assembly primers were designed using NEBuilder Assembly Tool (http://nebuilder.neb.com/), and primer sequence in lowercase mark the homology arm.

A 5.4kb region was identified around the *DDX5* promoter to be positive for the active chromatin marks H3K27ac and H3K4me3 (Wang et al., 2020). This fragment also included the first the first intron and exon of *DDX5*. This entire region was PCR amplified using Platinum™ PCR SuperMix High Fidelity (Thermo Fisher Scientific), genomic DNA as template, and the following primers: DDX5-PromoterGibson-F: aacatatccagtcactatggACCTGGAGTCACTGCTTG; DDX5-Intron1Gibson-R: acccccatacCCTGTGGTGAAAAAAGGAC. The PCR product was gel purified and sequence verified by comparing it to the killifish genome using BLAST. To amplify the DDX5-GFP gene cassette we used the DDX5-GFP yeast expression vector we generated as a PCR template (reference Yiwen’s section). The following primers were used (omitting the first exon which was included in the promoter fragment): DDX5-GFPGibson-F: tcaccacaggGTATGGGGGTGGCCCCCC; DDX5-GFPGibson-R: tggatcatcatcgatggtacTTACTTGTACAGCTCGTCCATGCCG. The PCR product was gel-purified and sequence-verified. The pDestTol2CV vector was digested with KpnI-HF and SalI-HF (NEB), removing the eye marker expression cassette (crystallin:Venus). The linearized plasmid was gel-purified. All three fragments were assembled using NEBuilder HiFi (NEB) according to the manufacturer’s protocol. The transposase expression vector was obtained from the Tol2Kit (http://tol2kit.genetics.utah.edu/index.php/Main_Page). 20 pg of plasmid DNA and 30 pg of transposase mRNA were co-injected into one-cell stage killifish embryos. Injected embryos were maintained at 26℃ and hatched at 15 days post injection. Overexpression of DDX5 protein in the F0 generation caused growth retardation and defects, resulting in failure to obtain germline transmission and an F1 generation. Therefore, only chimeric F0 animals with GFP expression in the brain were used in this study.

### GFP antibody staining in brain tissue

Three 7-month old F0 *Tg(ddx5:DDX5-GFP)* animals (males) with GFP expression in the brain were used for GFP antibody staining. Fish were euthanized in 500mg/mL MS-222 and the brain tissue were dissected out and subjected to overnight fixation with 4% PFA at 4 ℃. Brain cryosections were prepared as previously described (Harel et al., 2012; Harel et al., 2009). Antibody staining was done on 10 μm brain cryosections according to published protocol with modifications (Ganz et al., 2010). Briefly, brain cryosections were bleached with 5% H_2_O_2_ and 0.1% Formamide in PBS. Before primary antibody incubation, samples were treated with Trueblack™ Lipofuscin Autofluorescence Quencher (Biotium) for 30 s and no detergent was used in all follow-up steps. Anti-GFP antibody (ab13970, Abcam) and the goat anti-chicken secondary antibody Alexa 647 (Thermo Fisher) were used as 1:500. Samples were mounted in ProLong™ Gold Antifade Mount and imaged under Perkin Elmer Ultraview spinning disk.

### Generation of killifish DDX5 antibodies

Custom made antibodies aging the killifish DDX5 protein (XP_015809881.1) were generated in rabbits and performed by Bethyl, using their standard protocol that peptide synthesis, conjugation of antigen to carrier, subcutaneous immunizations of 2 rabbits, followed by ELISA testing results and affinity purification. The following peptide (corresponding to the C-terminus of killifish DDX5) was used as an immunogen: DDX5-625: 625-PYPIPPPFPQ-634. Antibodies were used at a dilution of 10,000 for western blot, and 1:500 for immunofluorescence.

### Tissue extracts and western-blot for DDX5 antibodies

Three young (2.5 months old) and three old (5.5 months old) *Nothobranchius furzeri* (GRZ strain) male fish were sedated in 200mg/L of Tricain (Sigma-Aldrich, A5040), and then euthanized in 500mg/L of Tricain in system water. Dissections were carried out under a stereo binocular (Leica S9E) at room temperature. Brains were dissected, and whole brains were placed in cold, slightly modified Buffer A (see Mass-Spec section: 30 mM Tris-HCl pH = 7.5, 40 mM NaCl, 3 mM CaCl_2_, 3 mM MgCl_2_, 5% glycerol, 1% Triton X-100) with freshly added protease inhibitors: pepstatin A at a final concentration of 1µM (Abcam, ab141416) and N-Ethylmaleimide at a final concentration of 10mM (Sigma-Aldrich, E3876). Homogenization was performed in a conical eppendorf tube to allow full homogenization of brain tissue with 3mm-metal beads (Eldan Israel, Cat# RC55 420). Homogenization was carried out with mechanical disruption using TissueLyzer LT (QIAGEN, #85600) with a dedicated adaptor (QIAGEN, #69980) at 50Hz for 4 minutes.

No centrifugation was performed, and whole brain lysate were used to avoid loss of DDX5 protein. Protein concentration was measured with BCA (Sigma-Aldrich), according to manufacture instructions. Brain homogenate was then mixed with a preheated (95°C) Sample Buffer with Urea (SB+U) (Tris-HCl pH6.8 62.5mM, Glycerol 10%, SDS 2%, 0.00125% and Urea 8M) at a ratio of 1:6 (sample to SB+U), boiled at 95°C for 5 minutes and placed on ice.

Standard western blot was performed. Briefly, 23µg of protein was resolved using Novex WedgeWell 4-20% Tris-Glycine gel (Thermo Fisher XP04205BOX), at 220V for 45 minutes. Transfer to a Nitrocellulose membrane was performed using the iBlot 2 (Thermo Scientific, IB21001). Membranes were then stained with Ponceau Red (Sigma, P7170-1L), and imaged with regular white light. Membranes were then washed in TBST 0.1% (Tris-HCl 10mM pH 8.0, NaCl 150mM, Tween 0.1%) 3 times for 5 min to remove Ponceau staining, and blocked in 5% BSA (sigma-aldrich) for 1 hour at room temperature. Next, anti-DDX5 (1:1000, custom-made, this paper) was incubated in 5% BSA blocking solution at 4°C overnight. Following washes, membranes were incubated with HRP secondary antibodies. For DDX5, we used anti-rabbit HRP (Abcam, ab6721). Membranes were treated with EZ-ECL kit and chemiluminescence was detected using Fusion Pulse 6 (Vilber Lourmat).

### Immunostaining of DDX5 in killifish brain sections

Fish were euthanized with MS222 (500mg/L) for 10 min at room temperature, and then the brains of 4 young (3.5 months), and 4 old (7 months) male fish were dissected immediately into cold PBS (pH 7.4, Gibco). The dissected brains were fixed with 4% paraformaldehyde (PFA) in PBS at 4°C overnight. The next day brain samples were transferred into 30% sucrose at 4°C overnight, followed by standard cryo-sectioning protocol. Samples were sectioned at 10 μM thickness and stained as previously described (Harel et al., 2012; Harel et al., 2009). Briefly, following three 10 min wash with PBST (PBS + 1% Tween), sections were blocked for 1h with serum-free block (Dako, X0909), and incubated over-night with our custom-made DDX5 antibodies (1:500) diluted in the blocking solution. Following subsequent washes with PBST, sections were incubated for 1 hr at room temperature with 1:300 goat anti-rabbit Alexa 488 secondary antibody (Abcam, ab150077)) and 2 μg/mL Hoechst. Samples were mounted in ProLong™ Gold Antifade Mount and imaged under Nikon Eclipse confocal microscope, using either X40 or X60 objectives.

### Immunostaining of DDX5 in mouse brain sections

All immunostainings were performed on C57BL/6 mice obtained from the NIA at the age indicated. Mice were subjected to intracardiac perfusion with 5mL of PBS containing heparin followed by 20mL of 4% paraformaldehyde (PFA, Electron Microscopy Sciences, RT 15714) in PBS. Brains were post-fixed for 24 hours in 4% PFA. They were then subjected to dehydration in 30% sucrose (Sigma-Aldrich, S3929) for 72 hours. Brains were subsequently embedded in Tissue-Tek optimal cutting temperature (O.C.T.) compound (Electron Microscopy Sciences, 62550) and sectioned using a cryostat in 12μm coronal sections that were mounted on glass slides. To perform immunofluorescent staining, sections were first washed with PBS, followed by permeabilization with ice cold methanol with 0.2% Triton-X for 10 min at room temperature. Sections were blocked with 5% Normal Donkey Serum (ImmunoReagents Inc., SP-072-VX10) and 1% BSA (Sigma Aldrich A8806) in PBS for 30 min at room temperature. Primary antibody staining was performed overnight at 4°C in 5% Normal Donkey Serum and 1% BSA in PBS. Primary antibodies used were the following: DDX5 (Atlas Antibodies, HPA020043, [1:200]) and NeuN (Millipore, MAB377, clone A60 [1:500]). Sections were washed with PBS with 0.2% Tween and then with PBS three times for 5 min at RT. Secondary antibody staining was performed at room temperature for 2 hr in 5% Normal Donkey Serum and 1% BSA in PBS. Secondary antibodies used were the following: donkey anti rabbit-AF568 (ThermoFisher [1:500]) and donkey anti mouse-AF488 (ThermoFisher [1:500]). Sections were washed with 0.2% Tween and then with PBS only three times for 5 min at room temperature. Sections were stained with DAPI (ThermoFisher 62248) at 1μg/mL for 10 min at room temperature. Sections were mounted with ProLong Gold (ThermoFisher, P36930) and visualized with a Nikon Eclipse Ti confocal microscope equipped with a Zyla sCMOS camera (Andor) and NIS-Elements software (AR 4.30.02, 64-bit) using the 60x objective. No blinding was done for picture taking. For visualization of images displayed in this manuscript, brightness and contrast were adjusted in Fiji to enhance visualization. The same settings were applied to all images shown for each experiment.

### Cloning of killifish cDNAs

Brain, liver, and muscle tissues from male African turquoise killifish *N. furzeri* were individually homogenized in RLT buffer (RNeasy Kit, # 74104 QIAGEN) using 0.5 mm Silica disruption beads (RPI-9834) and a tissue homogenizer (FastPrep-24, 116004500 - MP Biomedicals). Total RNA was isolated from the lysed tissues according to the RNeasy Kit protocol. cDNA for DDX5 was prepared from the total RNA from pooled brain, liver, and muscle tissues with high-capacity cDNA RT kit (Applied Biosystems, 4368814) using random primers, and according to the manufacturers’ protocol. cDNA was amplified using custom DNA oligonucleotides (IDT) and AccuPrime Pfx DNA Polymerase (Thermo Fisher Scientific). The following primers were used to clone killifish DDX5: DDX5-F: ATGCCTGGATTTGCAGACAG; DDX5-R: TTATTGTGGAAACGGTGGTG. PCR products were gel-purified, cloned using Zero Blunt TOPO PCR Cloning Kit (Thermo Fisher Scientific), and sequence-verified. Similarly, selected proteins with statistically significant increase in aggregation propensity in old brains compared to the young brains (HNRNPH1, MARK4, PABPN1, SCARB2) were cloned from the pooled cDNA (brain, liver, and muscle) using sequence-specific primers (available in Table S4C) and sequence-verified.

### Cloning and exogenous expression of killifish proteins in 293T cells

The sequence-verified ORFs of the full length and truncated versions of killifish DDX5 (1-483 [ΔIDR] or 1-535 [ΔPrD]) were cloned into pcDNA3.1 plasmid and transfected into human embryonic kidney 293T cells (HEK293T, ATCC, CRL-11268). The HEK293T cell line was not authenticated in-house. Mycoplasma testing was conducted every 3 months. The day before transfection, 9 × 10^6^ HEK293T cells were plated in a 10-cm dish in HEK293T medium (Dulbecco’s modified Eagle medium (DMEM, Invitrogen, 11965-092) supplemented with 10% fetal bovine serum (FBS Gibco, 16000-044), 1% 1% penicillin–streptomycin–glutamine (PSQ) (Gibco, 10378). The next day, the cells were transfected as follows: 100 μL of 1 mg/mL polyethylenimine (PEI; Polysciences, 23966-2, linear 25 kDa) was added to 2 mL of DMEM and incubated for 10 min at room temperature. The vector of interest (20 μg) was added to the PEI–DMEM mixture and incubated for 15 min at room temperature. The PEI–DMEM–DNA mixture was then added dropwise to the HEK293T cells and the medium was replaced with 8 ml fresh HEK293T medium 12h after transfection. Cell were imaged 24-36h after transfection.

### Cloning of DDX5 wild-type and mutants in yeast vectors

Using the DDX5 fragment we cloned from killifish cDNA, we designed primers to amplify either the region spanning amino acid residues 1-483 (ΔIDR) or 1-535 (ΔPrD). We then cloned the truncated mutants into a gateway entry vector pDONR221 (Invitrogen Cat# 12536017). We verified the sequence by sequencing and subcloned it into a yeast gateway vector to allow GAL-inducible expression of the ORF with a C-terminal EGFP or mRuby3 tag (pAG416GAL-ccdB-EGFP in Lindquist Advanced Gateway Vector Collection, (Alberti et al., 2007) or pAG426GAL-ccdB-mRuby3 generated for this paper) or a yeast gateway vector to allow constitutive expression of the mutant with a C-terminal EGFP tag (pAG415GPD-ccdB-EGFP in Lindquist Advanced Gateway Vector Collection, (Alberti et al., 2007)) in yeast.

### Exogenous expression of killifish proteins in *Saccharomyces cerevisiae* (one color overexpression assay)

The sequence-verified ORFs were then cloned into yeast gateway vector for GAL-inducible yeast expression with a fluorescent C-terminal EGFP tag (pAG416GAL-ccdB-EGFP in Lindquist Advanced Gateway Vector Collection, (Alberti et al., 2007)). The yeast expression plasmid (all low-copy centromeric-based) for each protein was transformed into BY4741 respectively and successful transformation was confirmed through prototrophy selection. The strains bearing the plasmids were inoculated overnight in 2% raffinose media (0.77 g of CSM-URA (Sunrise # 1004-010), 6.7 g of yeast nitrogen base without amino acid (BD 291920), and 20 g raffinose (RPI, # R20500-500.0) in 1 L media), then washed, diluted and switched to 2% galactose media (0.77 g of CSM-URA (Sunrise # 1004-010), 6.7 g of yeast nitrogen base without amino acid (BD 291920), and 20 g galactose (Fisher BioReagents, # BP656-500) in 1 L media) to induce protein expression. The overnight culture generally reached an OD_600_ of 0.9-1 and was further diluted back to OD_600_ ∼0.1 and induced for 6-8 hrs where the OD_600_ reach mid-log ∼0.4-0.6. We typically aliquot 2 µL of the mid-log phase yeast onto an imaging spot of a multi-spot microscope slide (6 mm diameter cell Thermo Cat# 9991090), mounted with cover glass (thickness 1 from Corning, Cat# 2975-246) and sealed with nail polish. Microscopy images were taken using a Leica inverted fluorescence microscope with a Hammamatsu Orca 4.0 camera. Exposure times in the DIC channel was 100ms and 50, 250, 500, and 1000 ms in the GFP channel (GFP excitation: 450–490 nm; emission: 500–550 nm; software: LASX DMI6000B; refraction index: 1.518; aperture: 1.4).

### Exogenous expression of killifish proteins in *Saccharomyces cerevisiae* (two-color seeding assay)

The full-length African killifish DDX5 and other proteins (PAPBN1, SNRNP70, ELAVL2, CK2A, Human DDX5, PolyQ, NONO, MAGI2, MARK4) were cloned into two yeast advanced gateway destination vectors: one with a constitutive GPD-promoter and a C-terminal EGFP-tag low copy number plasmid (p415GPD-ccdB-EGFP), another one with a galactose-inducible GAL-promoter and a C-terminal mRuby3-tag high copy number plasmid (p426GAL-ccdB-mRuby3). The two plasmids were co-transformed into BY4741 following standard lithium-acetate protocol. First, cells were inoculated and grown to saturation in rich media (YPD). The saturated cultures were then diluted by 10-fold and grew to mid-log. The cells were harvested, washed in sterile water twice, and resuspended in a transformation master mix (240 μL of PEG 3500 50% w/v, 36 μL1 M LiOAc, 50 μL boiled salmon sperm carrier DNA (2mg/mL), 14 μL plasmid DNA (0.1-1 μg), and sterile water to a final volume of 360 μL). Cells were incubated in the transformation master mix at 42°C for 45 min. Following the heat shock, cells were pelleted, and resuspended in 100 μL of sterile water and plated on respective selective medium.

The BY4741 strain co-transformed with the two fusion protein plasmids was first grown overnight to stationary phase in SD-URA-LEU. Images of the overnight culture were taken as pre-induction phase. The strains were then washed with 1x PBS three times and diluted to OD_600_ = 0.1 in SGAL-URA-LEU (2% galactose). The induction took approximately 6-10 h depending on when the culture reached mid-log (OD_600_ ∼ 0.4) and images were taken immediately after this induction phase. After induction, the strains were diluted by 200-fold (pipet 1 uL of SGAL-URA-LEU induction phase culture into 199 uL of SD-URA-LEU) SD-URA-LEU (2% glucose) and imaged after 24 h of growth as the outgrowth phase. We typically aliquoted 2 µL of the mid-log phase yeast onto an imaging spot of a multi-spot microscope slide (6 mm diameter cell Thermo Cat# 9991090), mounted with cover glass (thickness 1 from Corning, Cat# 2975-246) and sealed with nail polish. Microscopy images were all taken using a Leica inverted fluorescence microscope with a Hammamatsu Orca 4.0 camera. Exposure times in the DIC channel was 100ms and in the fluorescent channel around 50-500 ms (GFP excitation: 450–490 nm and emission: 500–550 nm; Cy3 excitation: and emission; software: LASX DMI6000B; refraction index: 1.518; aperture: 1.4; exposure time: 50, 250, and 500ms for both fluorescent channels).

The microscopy images were quantified using customized CellProfiler script (available in GitHub https://github.com/ywrchen/killifish-aging-aggregates) followed by extensive visual inspection and independent verification. We used the CellProfiler output to primarily identify and quantify total number of cells as well as total number of cells with positive GFP or Cy3 signals. Because the proteins we tested have heterogeneity in protein expression and GFP/Cy3 morphology, we used the CellProfiler foci scoring output as a reference (unified scoring code was difficult to implement) and counted all fluorescent images manually followed by blind counting done by two additional independent lab members on 20% of randomly selected raw images. The reported percentage of cells with foci was quantified as number of cells with GFP or Cy3 foci divided total number of cells with positive GFP or Cy3 signals. We only scored a cell as ‘foci-positive’ if it either had at least one focus along with diffuse GFP or Cy3 signals or had at least two foci inside a single cell. We considered each half of a budded yeast cells as two cells in the quantification.

### Fluorescence recovery after photobleaching

The yeast samples used for the fluorescence recovery after photobleaching (FRAP) experiment were prepared following the same procedure as the yeast one-color overexpression assay described above. Briefly, laboratory strain BY4741 transformed with pA416GAL-DDX5(killifish)-EGFP and [*RNQ*^+^][*psi*^-^] strain transformed with pAG416GAL-Sup35NM(yeast)-EGFP were inoculated in 1 mL of 2% raffinose media (0.77 g of CSM-URA (Sunrise # 1004-010), 6.7 g of yeast nitrogen base without amino acid (BD 291920), and 20 g raffinose (RPI, # R20500-500.0) in 1 L media). The next day, the overnight culture was spun down (3000 g x 5 min at room temperature), washed with 1X PBS twice, diluted by ∼10 fold (typical OD600 for overnight culture can reach 0.8-1 and the dilution aimed to dilute the culture to OD600 ∼0.1) and switched to 1 mL of 2% galactose media (0.77 g of CSM-URA (Sunrise # 1004-010), 6.7 g of yeast nitrogen base without amino acid (BD 291920), and 20 g galactose (Fisher BioReagents, # BP656-500) in 1 L media) to induce protein expression. We typically waited 6-8 hrs, allowing the culture to undergo 2-3 rounds of doubling (mid-log OD_600_ ∼0.4-0.6) for sufficient protein induction before imaging.

We prepared thin agar pad for the samples where 8 µL of semi-cooled 1.5% agarose solution (add low melting temperature NuSieve GTG Agarose (Lonza, Cat# 50081) to 2% galactose media (0.77 g of CSM-URA (Sunrise # 1004-010), 6.7 g of yeast nitrogen base without amino acid (BD, 291920), and 20 g galactose (Fisher BioReagents, # BP656-500) in 1 L media), heat it to ∼60°C and cool for 1 min) was loaded to each imaging spot of a multi-spot microscope slide (6 mm diameter cell Thermo Cat# 9991090). Typically, 2 µL of the mid-log phase yeast was loaded onto the freshly made agar pad, mounted with high performance cover glass #1.5 (0.17mm +/- 5.0 micron thick, Zeiss #474030-9000-000) and sealed with nail polish. Images were acquired through 3D Structured Illumination microscopy (SIM) with the DeltaVision OMX BLAZE system (Applied Precision-GE, Inc.) equipped with 3 emCCDs (Evolve, Photometrics Inc.) using an Olympus UPlanApo 100x (NA 1.40) oil-immersion objective. SIM excitation was with 100 mW lasers (405nm 488nm, 568nm), widefield epifluorescence excitation for deconvolution datasets was with InsightSSI™ illuminator (405nm, 488nm, 568 nm) and standard emission filter sets (528/48 nm and 609/37 nm). We used DeltaVision OMX’s built-in FRAP and photo kinetic module to set up the time-series experiment. Before each photobleaching experiment, we first drew rectangles around the desired photobleaching areas and recorded their sizes and coordinates. We acquired 3 frames before the actual photobleaching event at the same exposure (FITC-GFP in emCCD-mode with a gain of 500 at 100% intensity for total 100 ms excitation time) and interval (every 20 s) as the recovery stage. The single photobleaching event occurred at frame #4 in the 488-laser channel (GFP) at 31.3% intensity for 0.1 seconds. We continuously imaged throughout the fluorescence recovery stage for a total of 5 minutes at 20 seconds interval.

We analyzed the FRAP time-series images in Fiji. The images were exported as virtual stacks and went through a registration step to minimize drift that would contribute to signal fluctuation. Registration was done with Fiji plugins RegStack or TurboStack followed by manual inspection and the resulting image stacks were used for downstream analysis. Next, we drew the exact photobleaching area (with location and size information recorded at the image acquisition time) and recorded them as region of interests (ROIs). We drew 2X additional areas (ROIs) that were the same size as the photobleached areas in the background as well as inside cells where no photobleaching event occurred. Last, the intensity values of all these ROIs including minimum, mean, and maximum intensity values were exported using the ROI Manager in Fiji. We used the average intensity values of the image background as baseline and subtract that from the photobleached as well as un-bleached ROIs. The FRAP curve was constructed from all photobleached areas after background subtraction (done for each timepoint) and the average as well as standard error of mean were shown. All images for the FRAP experiments were acquired at Stanford Cell Sciences Imaging Facility.

### Required Acknowledgement

The project described was supported in part by Award Number 1S10OD01227601 from the National Center for Research Resources (NCRR). Its contents are solely the responsibility of the authors and do not necessarily represent the official views of the NCRR or the National Institutes of Health.

### DDX5-SNAP *E. coli* expression and purification

The full-length African killifish DDX5 was cloned into the pDEST17 vector containing an N-terminal His10-Smt3-tag and a C-terminal SNAP-tag (New England BioLabs). *E. coli* LOBSTR-BL21(DE)-RIL (kerafast) host cells were transformed with the killifish DDX5 plasmid and were grown to an OD of ∼0.4, induced by 0.1 mM IPTG overnight at 17 °C in LB medium supplemented with 100 µg/mL ampicillin and 25 µg/mL chloramphenicol. Cells were pelleted and resuspended in lysis buffer (50 mM Tris-HCl pH 7.4, 250 mM NaCl, 10% (vol/vol) glycerol, 1 mM DTT, and 10 mM imidazole) containing 200 µg/mL lysozyme and an EDTA-free protease inhibitor cocktail (Roche Diagnostics, cOmplete^™^, EDTA-free Protease Inhibitor Cocktail). After 30 min of lysozyme digestion at 4 °C with constant mixing, cells were lysed by sonication, and cellular debris was pelleted at 30,000 g for 30 min. Cleared lysate was incubated with Ni-NTA agarose resin (Qiagen), washed well with Ni-Wash buffer (50 mM Tris-HCl, pH 7.5, 500 mM NaCl, 1 mM DTT, 10% (vol/vol) glycerol, and 25 mM imidazole), and eluted with Ni-Elution buffer (50 mM Tris-HCl, pH 7.5, 500 mM NaCl, 1 mM DTT, 10% (vol/vol) glycerol, and 200 mM imidazole). The eluted protein was dialyzed in high salt storage buffer (50 mM Tris-HCl, pH 7.5, 1 M NaCl, 10% (vol/vol) glycerol, and 1 mM DTT) and digested overnight with 2 µg/mL His6-Ulp1 (Smt3/SUMO-specific protease, purified from *E. coli* with nickel affinity resin) at 4 °C to remove the N-terminal His10-SMT3 tag. The digested protein was incubated with Ni-NTA agarose resin again for a second round of native affinity purification, washed well with Ni-Wash high salt buffer (50 mM Tris-HCl, pH 7.5, 1 M NaCl, 1 mM DTT, and 10% (vol/vol) glycerol), and eluted with Ni-Elution high salt buffer (50 mM Tris-HCl, pH 7.5, 1 M NaCl, 1 mM DTT, 10% (vol/vol) glycerol, and 25 mM imidazole). We performed size exclusion chromatography on the eluted proteins in a Superdex 200 Increase 10/300 GL column (GE Healthcare) using an Akta Ettan fast protein liquid chromatography (FPLC) system (GE Healthcare) to further separate the desired protein from contaminants. The resulting protein was concentrated (Amicon Ultra 0.5 mL and 4 mL Centrifuge filters, 30 kDa NMWL devices, Millipore Cat#UFC505024 and UFC805024) and flash frozen in liquid nitrogen for future use.

### DDX5(1-535)-SNAP *E. coli* expression and purification

The DDX5 mutant 1-535 lacking the putative C-terminal prion-like domain (PrD) was cloned into the pDEST17 vector containing an N-terminal His10-Smt3-tag and a C-terminal SNAP-tag (New England BioLabs). The same purification scheme was used to purify DDX5(1-535)-SNAP than DDX5-SNAP, except the native affinity purification were done with medium salt concentration and the protein was flash-frozen in medium salt storage buffer (50 mM Tris-HCl, pH 7.5, 500 mM NaCl, 1 mM DTT, and 10% (vol/vol) glycerol).

### DDX5(1-483)-SNAP *E. coli* expression and purification

The DDX5 mutant 1-483 lacking the intrinsically disordered region (IDR) was cloned into pDEST17 vector containing an N-terminal His10-Smt3-tag and a C-terminal SNAP-tag (New England BioLabs). The exact same purification scheme for DDX5(1-535)-SNAP was used to purify DDX5(1-483)-SNAP and the protein was flash frozen in medium salt storage buffer (50 mM Tris-HCl, pH 7.5, 500 mM NaCl, 1 mM DTT, and 10% (vol/vol) glycerol).

### His10-DDX5-SNAP *E. coli* expression and purification

The full-length African killifish DDX5 was cloned into the pDEST17 vector containing an N-terminal His10-tag and a C-terminal SNAP-tag (New England BioLabs). *E. coli* LOBSTR-BL21(DE)-RIL (kerafast) host cells were transformed with the killifish DDX5 plasmid and were grown to an OD of ∼0.4, induced by 0.1 mM IPTG overnight at 17 °C in LB medium supplemented with 100 µg/mL ampicillin and 25 µg/mL chloramphenicol. Cells were pelleted and resuspended in lysis buffer (50 mM Tris-HCl pH 7.4, 250 mM NaCl, 10% (vol/vol) glycerol, and 1 mM DTT) containing 200 µg/mL lysozyme and an EDTA-free protease inhibitor cocktail (Roche Diagnostics, cOmplete^™^, EDTA-free Protease Inhibitor Cocktail). Keep a 1:20 lysis buffer to culture volume ratio. After 30 min of lysozyme digestion at 4 °C with constant mixing, cells were lysed by sonication (five 2 min cycle, 2s on 2s off, at 100% amplitude, all performed on ice), and cellular debris was pelleted at 30,000 x g (Beckman JA25.25 rotor) for 30 min. Cleared lysate was incubated with Ni-NTA agarose resin (Qiagen), washed well with Ni-Wash buffer (50 mM Tris-HCl, pH 7.5, 200 mM NaCl, 0.5 mM DTT, and 10% (vol/vol) glycerol), and eluted with Ni-Elution buffer (50 mM Tris-HCl, pH 7.5, 200 mM NaCl, 0.5 mM DTT, 10% (vol/vol) glycerol, and 200 mM imidazole). The eluted protein was dialyzed at 4 °C to remove imidazole (dialysis buffer: 50 mM Tris-HCl, pH 7.5, 200 mM NaCl, 200 mM arginine, 10% (vol/vol) glycerol, and 0.5 mM DTT) and concentrated to ∼ 5 mg/mL (Amicon Ultra 4 mL Centrifuge filters, 30 KDa NMWL devices, Millipore Cat# UFC805024). Last, we performed size exclusion chromatography on the concentrated proteins in a Superdex 200 Increase 10/300 GL column (GE Healthcare) using an Akta Ettan fast protein liquid chromatography (FPLC) system (GE Healthcare) to further separate the desired protein from contaminants. The resulting protein was further concentrated (Amicon Ultra 0.5 mL and 4 mL Centrifuge filters, 30 KDa NMWL devices, Millipore Cat#UFC505024 and UFC805024) and flash frozen in liquid nitrogen for future use.

### Protein labeling reaction

Recombinant proteins were labeled with SNAP-Surface 549 dye (New England BioLabs, Cat# S9112S) for visualization. The labeling reaction was performed in the high salt storage buffer (for DDX5-SNAP) or medium salt storage buffer (for DDX5(1-535)-SNAP and DDX5(1-483)-SNAP) described above while keeping the SNAP-Surface dye at 2x the final concentration of the protein. To minimize protein degradation and photobleaching of the dye, we followed the following order of the reagent addition was: 1) recombinant protein, 2) 4x storage buffer, 3) 5% Tween-20, 4) 100 mM DTT, and 5) 2x SNAP Surface 549 dye to minimize proteolysis. The labeling reaction was incubated overnight in dark (to avoid photobleaching) at 4°C. To remove excess non-covalently linked SNAP dye, we used a standard desalting column (Zeba spin desalting columns pre-equilibrated with high salt buffer (1 M NaCl, 50 mM Tris-HCl, pH 7.5, 1 mM DTT for DDX5-SNAP) or medium salt buffer (500 mM NaCl, 50 mM Tris-HCl, pH 7.5, 1 mM DTT for DDX5(1-535)-SNAP and DDX5(1-483)-SNAP), Thermo Fisher, Cat# 89882). In case of preparing purified recombinant protein inputs for the ATPase activity assay, we used dialysis exclusively to remove excess dye while adjusting the salt concentration to a desired level (50 mM NaCl) at the same time (Slide-A-Lyzer^TM^ 20K MWCO Mini dialysis cassette were used, Thermo Cat#69590). Successful labeling was verified by running SDS-PAGE and image the gel directly using Bio-Rad Universal Hood II Gel Doc Molecular Imaging System CFW-1212M (using the protein blot DyLight 550 setting, excitation/emission: 550/518). We ran a small amount (1 µL) of Bio-Rad Precision Plus Protein WesternC standards along with the labeled protein while keeping at least one open lane in between the protein ladder and sample to avoid signal bleed-through. We imaged immediately after gel electrophoresis to obtain the best signal.

### Phase diagrams

Purified proteins were labeled overnight (the same protein storage buffer with additional 2x dye for every 1x protein, 1mM DTT, 5% Tween-20) with SNAP DyLight 549 dye. After labeling, proteins were resolved on SDS-PAGE and imaged on a gel doc system with excitation/emission at 550/518nm (see protein labeling reaction section for details) to ensure the correct protein was labeled and there was minimum degradation. Phase diagrams were generated through dialysis using Slide-A-Lyzer MINI Dialysis Device (20K MWCO, 0.1 mL, Thermo Cat#69590) and the resulting protein was further diluted to respective protein/salt mixture (no inclusion of glycerol, DNA, RNA, or ATP in any condition) when necessary. The protein concentration for each condition was measured directly from 1 µL aliquot of the sample using NanoDrop (record A280nm measurements and use the exact same protein sample buffer as blank) and the Bradford assay (Biorad). After incubation for different amounts of time (1 hour after dialysis or dilution to allow droplet maturation) at different temperatures (either 4°C or 25°C), 1-2 µL of the protein/salt mixtures was aliquoted onto a coverslip/glass slide (multi-spot microscope slide (6 mm diameter cell Thermo Cat# 9991090) mounted with cover glass (thickness 1 from Corning, Cat# 2975-246)) and sealed with nail polish. Microscopy images were taken using a Leica inverted fluorescence microscope with a Hammamatsu Orca 4.0 camera. Exposure times in the DIC channel was 100ms and in the fluorescent channel around 50-500 ms (GFP excitation: 450–490 nm and emission: 500–550 nm; Cy3 excitation: and emission; software: LASX DMI6000B; refraction index: 1.518; aperture: 1.4; exposure time: 50, 250, and 500ms for both fluorescent channels). Scoring for droplet formation was based on DIC and Cy3 imaging using a 100x oil objective on an inverted Leica microscope. If spherical-shaped droplets were observed in both DIC and Cy3 channel, we noted that query protein can phase separate under the particular protein and salt concentrations. We used results conducted at 4°C to construct the phase diagram.

### Salt, protein concentration, temperature, and time effects on DDX5 assembly

We assessed the spontaneous assembly of DDX5 at different salt and protein concentrations at different temperatures in the absence of nucleic acids (DNA or RNA) and ATP. We focused on four conditions where DDX5 was shown to phase separate. Specifically, full length DDX5 stock protein (∼24 µM stored in 1 M NaCl, 50 mM Tris-HCl, pH 7.4, 10% glycerol, 1 mM DTT at −80°C) was labeled overnight at 4°C (see labeling reaction for detail) with SNAP DyLight 549 dye. After labeling, proteins were resolved on SDS-PAGE and imaged on a gel doc system with excitation/emission at 550/518nm (see protein labeling reaction section for details) to ensure the correct protein was labeled and there was minimum degradation. Next, we dialyzed 20 µL of purified proteins using Slide-A-Lyzer MINI Dialysis Device (20K MWCO, 0.1 mL) in 250 mL medium salt buffer (500 mM NaCl, 50 mM Tris-HCl, pH 7.4, 1 mM DTT) and 250 mL low salt buffer (150 mM NaCl, 50 mM Tris-HCl, pH 7.4, 1 mM DTT), respectively, for a total of 1 hour with constant stirring (200 r.p.m.) at 4°C. After dialysis, protein concentration in either sample were estimated from nanodrop A280nm measurements (the exact same protein sample buffer was used as blank to zero the reads). We further diluted the proteins to the following 4 conditions (condition 1: [NaCl] = 500 mM and [DDX5] = 9.8 µM; condition 2: [NaCl] = 250 mM and [DDX5] = 4 µM; condition 3: [NaCl] = 150 mM and [DDX5] = 6 µM; condition 4: [NaCl] = 50 mM and [DDX5] = 2.8 µM all in 50 mM Tris-HCl, pH 7.4, 1 mM DTT, no RNA/DNA/ATP) and prepared 5 µL sample for each condition for incubation at either 4°C or room temperature (∼25°C) with no shaking/agitation. At 2 hr and 24hr timepoints, we aliquoted 1 µL of protein from each condition onto a coverslip (multi-spot microscope slide (6 mm diameter cell Thermo Cat# 9991090) mounted with cover glass (thickness 1 from Corning, Cat# 2975-246)), sealed with nail polish, and imaged them. Microscopy images were taken using a Leica inverted fluorescence microscope with a Hammamatsu Orca 4.0 camera using a 100x oil objective. Exposure times in the DIC channel was 100ms and in the fluorescent channel around 50-500 ms (GFP excitation: 450–490 nm and emission: 500–550 nm; Cy3 excitation: and emission; software: LASX DMI6000B; refraction index: 1.518; aperture: 1.4; exposure time: 50, 250, and 500ms for both fluorescent channels). The microscopy images of *in vitro* assembled protein condensate were quantified using CellProfiler. Specifically, we used a customized CellProfiler script (available in GitHub https://github.com/ywrchen/killifish-aging-aggregates) to identify and quantify the total number of condensates and characterize their shape (form factor) and intensity.

### In vitro seeding

Full length killifish DDX5 stock protein (∼24 µM of DDX5-SNAP stored in 1 M NaCl, 50 mM Tris-HCl, pH 7.4, 10% glycerol, 1 mM DTT at −80°C) was labeled overnight at 4°C (see labeling reaction for detail) with SNAP DyLight 549 dye. A killifish DDX5 variant with an N-terminal His10-tag (∼16 µM of His10-DDX5-SNAP stored in 1 M NaCl, 50 mM Tris-HCl, pH 7.4, 10% glycerol, 1 mM DTT at −80°C) was labeled overnight at 4°C (see labeling reaction for detail) with SNAP Alexa 488 dye. After labeling, proteins were resolved on SDS-PAGE and imaged on a gel doc system with excitation/emission at 550/518nm (see protein labeling reaction section for details) to ensure the correct protein was labeled and there was minimum degradation. The labeled DDX5-SNAP protein was subjected to desalting where the column (Zeba spin desalting columns, Thermo Fisher, Cat# 89882) was pre-equilibrated with high salt buffer (1 M NaCl, 50 mM Tris-HCl, pH 7.4, and 1 mM DTT) and the resulting protein was diffuse serving as “recipients” (final stock concentration was 4.35 µM determined by Bradford). The N-terminal His10-tagged DDX5-SNAP was dialyzed in low salt buffer (150 mM NaCl, 50 mM Tris-HCl, pH 7.4, and 1 mM DTT) to induce the formation of amorphous aggregates which served as the “seed” (final stock concentration was 8.7 µM determined by Bradford). We set up the *in vitro* assembly with equal volume of diffuse DyLight 549 labeled DDX5-SNAP and equal volume of three 5-fold serial dilutions (diluent was the same low salt buffer) of Alexa 488 labeled His10-DDX-SNAP aggregates in the absence of DNA/RNA/ATP. We also included a control where the diffuse DyLight 549 labeled DDX5-SNAP was mixed with equal volume of low salt buffer only. All samples were incubated at room temperature (∼25°C) with no shaking/agitation. At 1 hr, 24hr, and 72 hour timepoints, we aliquoted 1 µL of protein from each condition onto a coverslip (multi-spot microscope slide (6 mm diameter cell Thermo Cat# 9991090) mounted with cover glass (thickness 1 from Corning, Cat# 2975-246)), sealed with nail polish, and imaged them. Microscopy images were all taken using a Leica inverted fluorescence microscope with a Hammamatsu Orca 4.0 camera using a 100x oil objective. Exposure times in the DIC channel was 100ms and in the fluorescent channel around 50-500 ms (GFP excitation: 450–490 nm and emission: 500–550 nm; Cy3 excitation: and emission; software: LASX DMI6000B; refraction index: 1.518; aperture: 1.4; exposure time: 50, 250, and 500ms for both fluorescent channels).

### ATPase assay

DDX5 is an ATP-dependent helicase (Hirling et al., 1989). To test the ATPase activity of DDX5, ATPase assays were set up in 96-well PCR plate on ice where fresh ATP stock was added last. A 10 µL reaction was set up with final condition of 50 mM Tris-HCl pH 7.5, 150 mM NaCl, 5 mM MgCl_2_, 2 mM ATP, variable amount of protein, and 500 ng/µL of yeast total RNA (unless otherwise noted). Reactions were conducted at room temperature for 30 minutes (or for variable amount of time specified in the time-course experiments), diluted to a final volume of 50 µL by addition of 40 µL nuclease-free water and quenched with 100 µL of BIOMOL GREEN (molybdate/malachite green-based colorimetric phosphate assay). The rate of ATP hydrolysis was reported as amount of free phosphate released in a unit time and volume. The free phosphate is inferred from OD620_nm_ reading upon 20 or 30 minutes of color development after BIOMOL GREEN addition. A phosphate standard curve is constructed beforehand in the exact reaction buffer condition to relate OD620_nm_ reading and the amount of free phosphate in solution. In all experiments, we set up >4 technical replicates for each condition. All experiments were independently repeated for at least three times and the average and standard error of mean were shown.

### Droplet and diffuse protein activity comparison

Full-length DDX5 or ΔPrD or ΔIDR mutants were labeled overnight at 4°C the night before use. To induce robust formation of droplets, full-length DDX5 or ΔPrD or ΔIDR mutants were subjected to dialysis in droplet induction buffer (50 mM NaCl, 50 mM Tris-HCl, 0.5 mM DTT) at 4°C for 15 minutes. Dialysis (a 100 µL dialysis cassette was used at maximum 40% volume capacity, Slide-A-Lyzer 0.1 mL MINI Dialysis Device 20K MWCO, Thermo Cat#69590) was done in 10000:1 buffer to sample volume ratio (or even larger) to ensure rapid equilibration of salt to the desired 50 mM concentration without drastic increase in sample volume. For a 30 µL aliquot, 15 min of dialysis was generally enough to equilibrate the salt in our setup, which included a stir bar and rotation speed at 200 r.p.m.. The dialyzed sample was imaged right away to confirm the presence of droplets. For samples that require droplet as starting material, the dialyzed sample served as the protein input. Diffuse sample was obtained by selecting the supernatant after a spin at 13,000 g for 20 min at 4°C for experiments described in Figure 5. For experiments described in Figure 6, the resulting pre-steady state diffuse sample were divided in half where one was further incubated at 4°C to allow enough time for new droplet to mature. This second experimental setup ensured that the total protein concentration of the diffuse and droplet samples remained the same. Protein concentration was estimated from nanodrop A280nm measurements for each condition (using the same protein sample buffer as blank to zero the reads). We did not observe that the presence of droplet upon thorough mixing significantly altered the nanodrop reading. The ATPase activity assay on droplet and diffuse protein was performed as described above in the ATPase reaction section.

## SUPPLEMENTAL FIGURE LEGENDS

**Figure S1 – Related to Figure 1.**
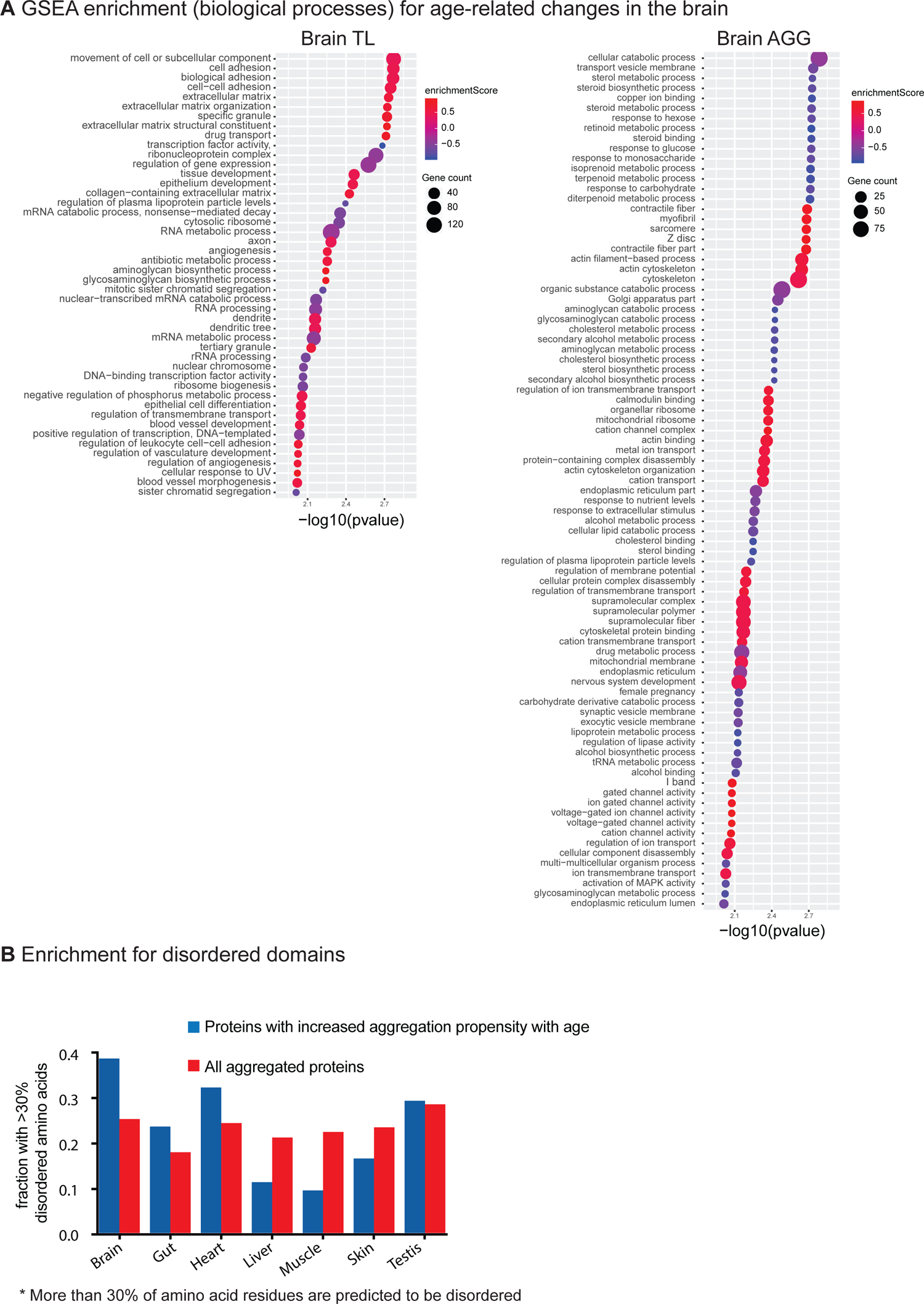
**(A)** GO enrichment using Gene Set Enrichment Analysis (GSEA) for proteins with increased age-related changes in the brain, for either tissue lysate (TL, left), or aggregate fraction (AGG, right). Enrichment score = score for enrichment/depletion of GO terms in this gene set. Counts = number of genes. **(B)** Enrichment for intrinsically disordered regions in proteins with increased aggregation propensity with age in the brain. Proteins with more than 30% of amino acid residues that are predicted to be disordered are considered here. Red: all detected aggregated proteins; Blue: proteins with an age-related increase in aggregation propensity are in blue. No significant difference between the two groups in each organ (Fisher’s exact test).

**Figure S2 – Related to Figure 2.**
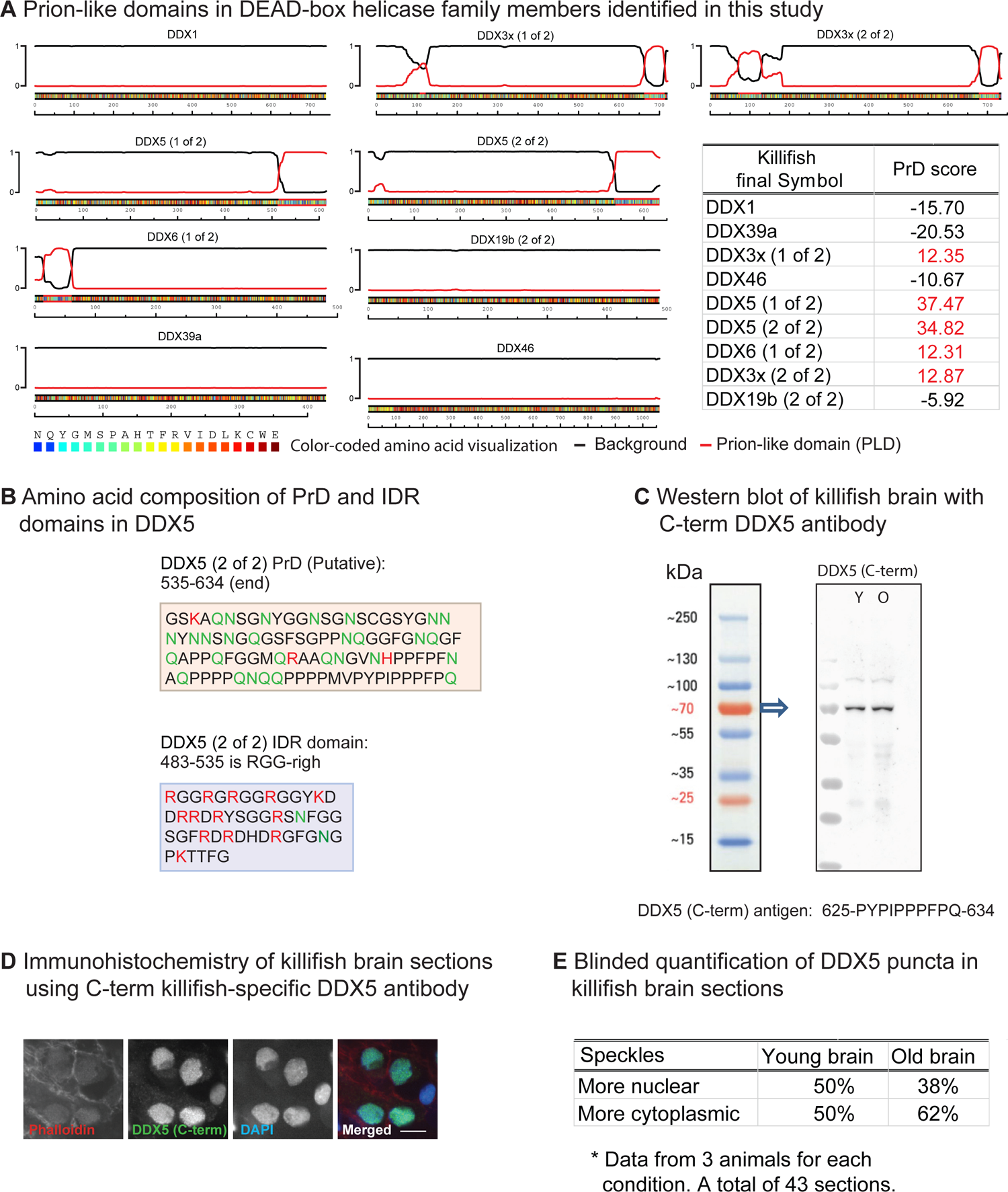
**(A)** Location of the predicted prion-like (PrD) domain in DEAD-box helicase family members identified in this study. Red line: score of the PrD domain. Amino-acid numbers are indicated at the bottom. PrD score (PLAAC) is indicated (bottom right). Amino-acid composition is color-coded. **(B)** Amino acid composition of PrD and RGG (motifs rich in arginines and glycines) domains in the primary DDX5 paralog in this study, DDX5 (2 of 2). The DDX5 (1 of 2) is more prevalent in the liver. Amino-acid numbers of each domain are indicated. **(C)** Production and validation of killifish DDX5 C-terminus antibody using a Western blot for brain lysates, extracted from either young (∼3 month old) or old (∼7 month old) killifish. The peptide sequence used as an antigen for immunization is indicated at the bottom. **(D)** Immunohistochemistry of killifish brain sections using C-term DDX5 antibody (green) and phalloidin, that recognizes staining actin filaments (red). Blue: DAPI. Scale bar: 10µm. **(E)** Qualitative quantification of the localization of DDX5 puncta in brain sections from young (3.5 months old) and old (7 months old) killifish. Data from 3 animals for each condition, with a total of 43 sections.

**Figure S3 – Related to Figure 2.**
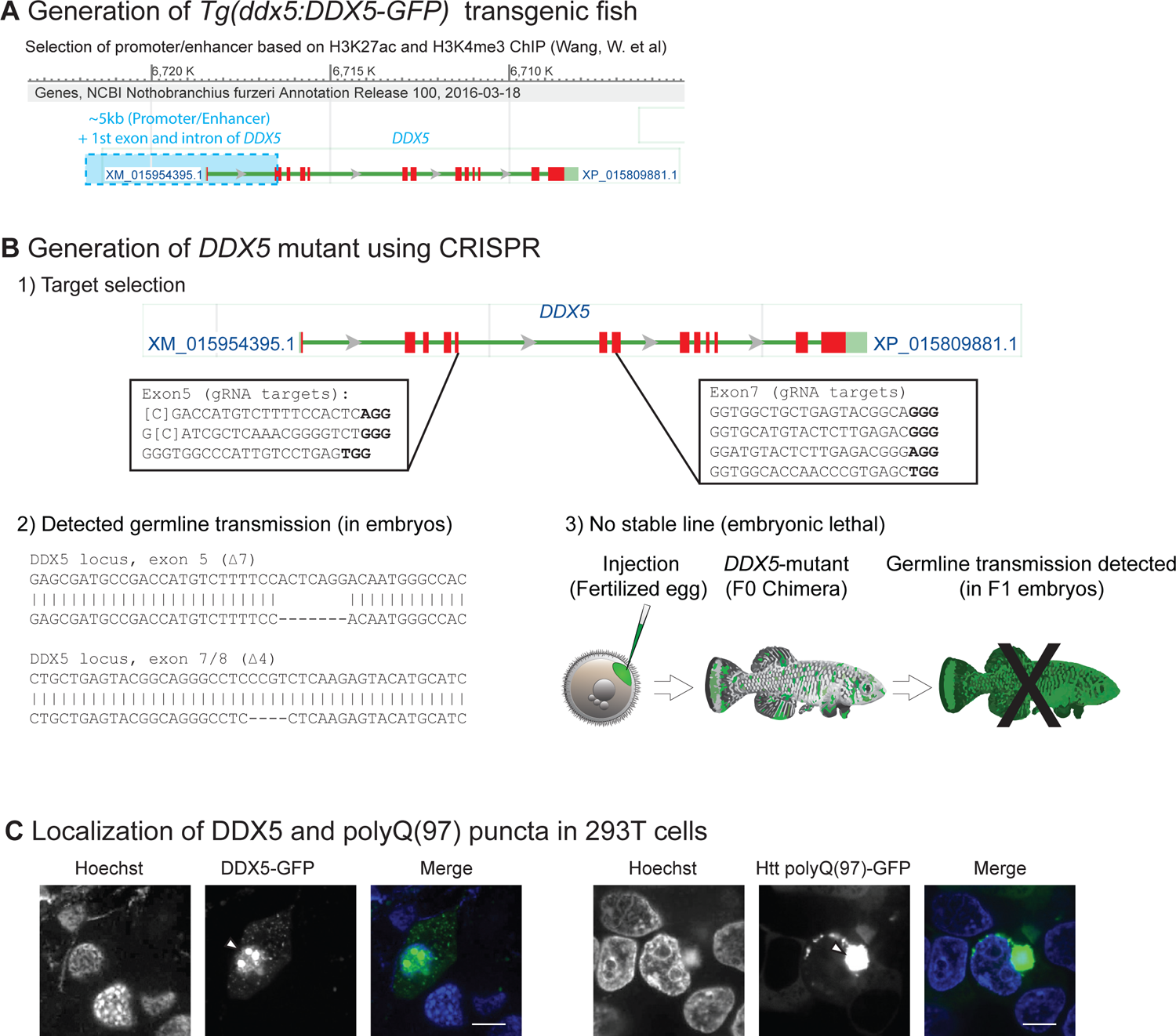
**(A)** The *Tg(ddx5:DDX5-GFP)* transgenic line was generated by using 5kb surrounding the transcriptional start site of the *DDX5* gene (including the first exon) to drive the expression of DDX5-GFP fusion protein (dashed blue line). Exons are indicated in red. **(B)** Generation of DDX5 CRISPR killifish mutant, including (1) gRNA target design for exons 5 and 7, and (2) evidence for successful germline transmission in F1 embryos. (3) F1 fries were never detected, even as heterozygous. Therefore, no stable line was generated. **(C)** Localization of DDX5-GFP (left) or polyQ(97)-GFP puncta in 293T cells. DDX5 puncta are primarily nuclear, whereas polyQ(97) puncta are primarily perinuclear (white arrowheads). Green: GFP-tag of the indicated protein; Blue: Hoechst. Images are representative of 2 experiments, each performed in triplicates. Scale bar: 5µm.

**Figure S4 – Related to Figure 3.**
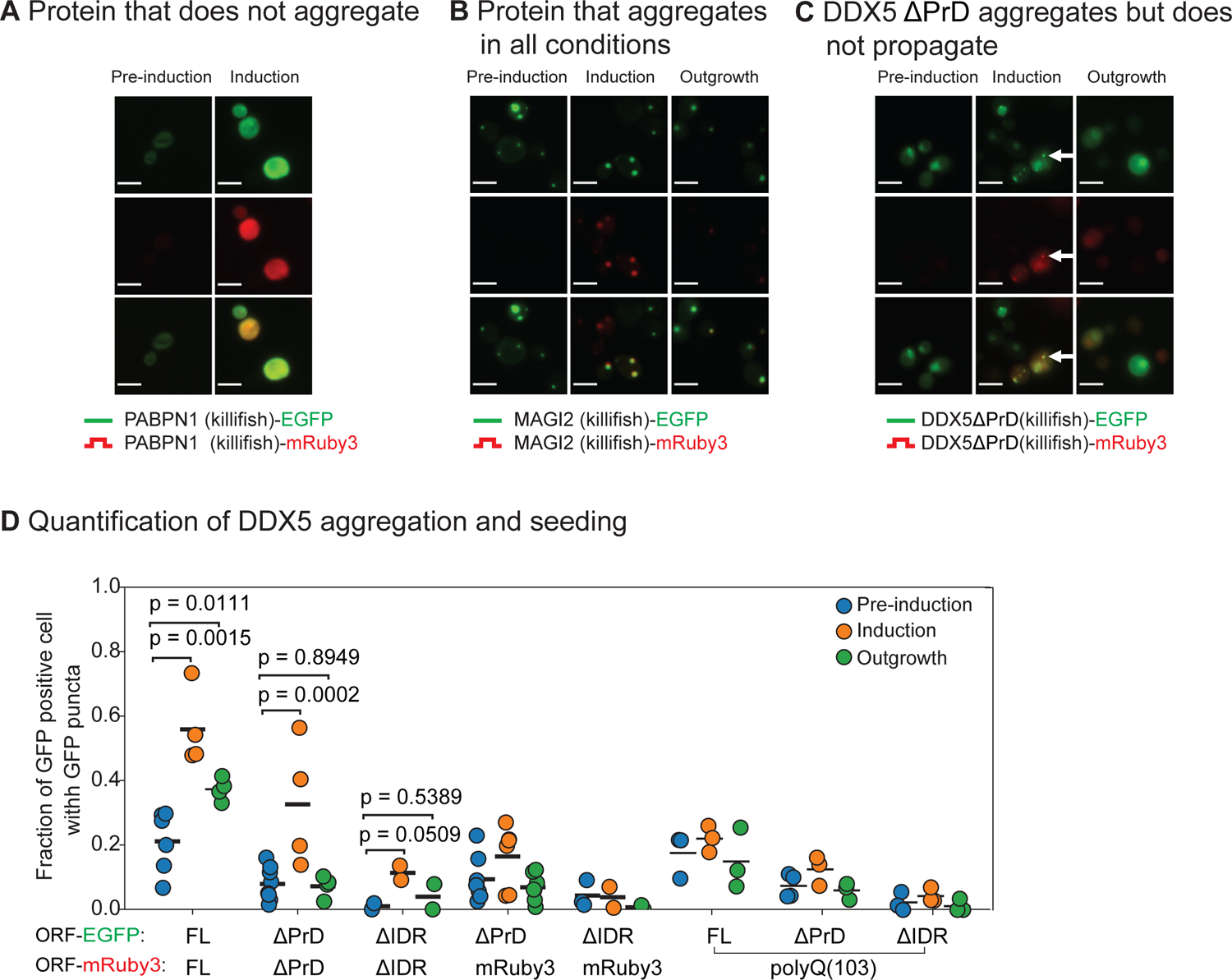
**(A, B)** Identification of killifish candidate proteins that do not aggregate (A), and proteins that aggregate under basal expression levels (B). **(C)** Following overexpression of DDX5ΔPrD-mRuby3, DDX5ΔPrD-GFP form aggregates (white arrows), which become diffuse and nuclear-localized after propagation. Representative of 4 independent experiments (quantified in D). Scale bar: 5µm. **(D)** Quantification of DDX5 aggregates and prion-like seeding potential by calculating the fraction of GFP positive cells with EGFP puncta in the indicated systems during pre-induction, induction, and outgrowth. Mean values from 3 or more independent experiments are indicated by the black bar, and each dot represents the average value within each experiment (induction and outgrowth phase results were selected when overexpression of mRuby3-tagged protein was robust. An average of 135 GFP-positive yeast cells are quantified for each strain under each experimental condition). P-values: from unpaired t-test between indicated experimental phases.

**Figure S5 – Related to Figure 5.**
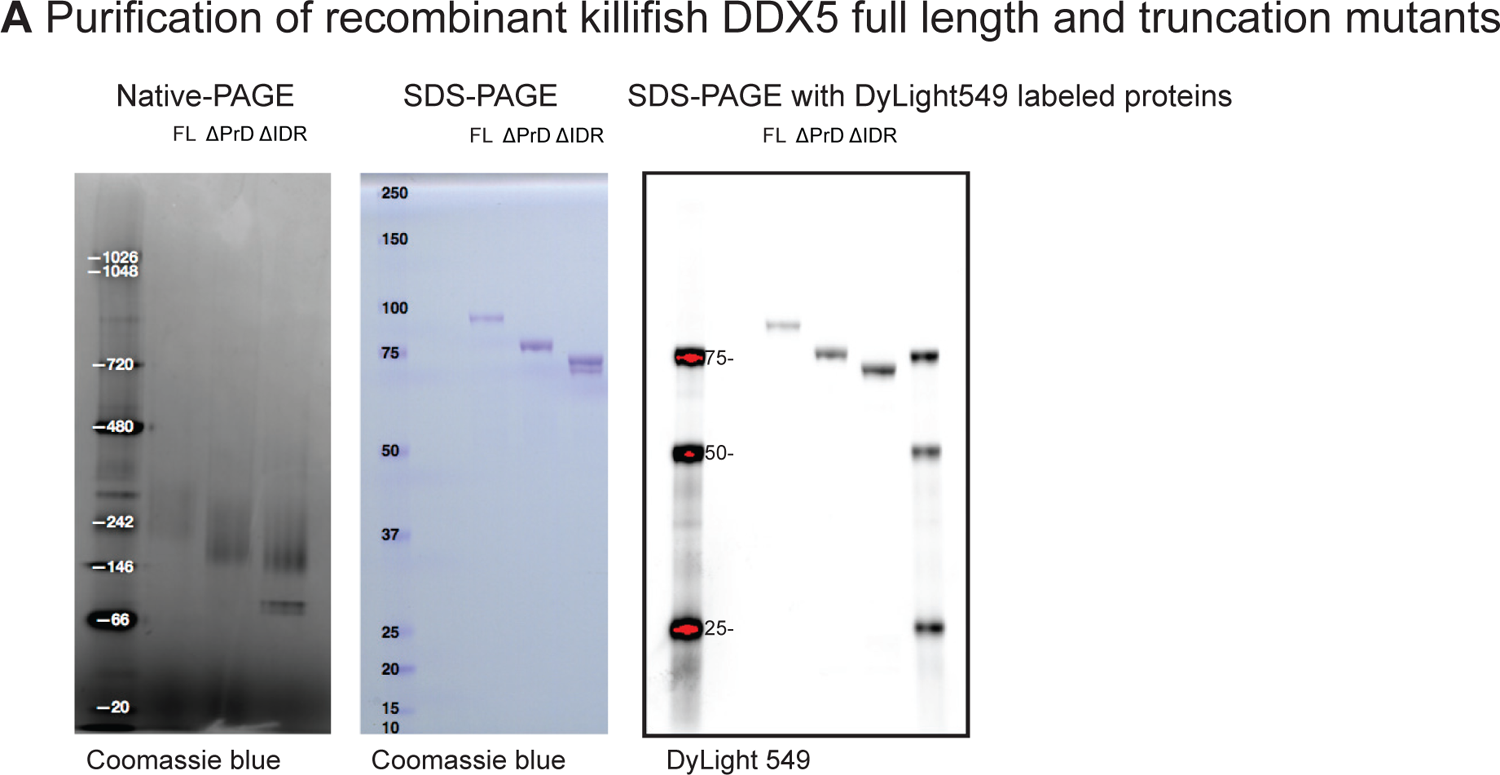
**(A)** Purified recombinant killifish DDX5 FL and mutants (ΔPrD and ΔIDR), as seen by Native-PAGE stained with coomassie blue (left), SDS-PAGE stained with coomassie blue (center), and SDS-PAGE blotted against DyLight 549 (SNAP-tagged protein was labeled with DyLight 549 dye prior to gel electrophoresis) (right).

**Figure S6 – Related to Figure 6.**
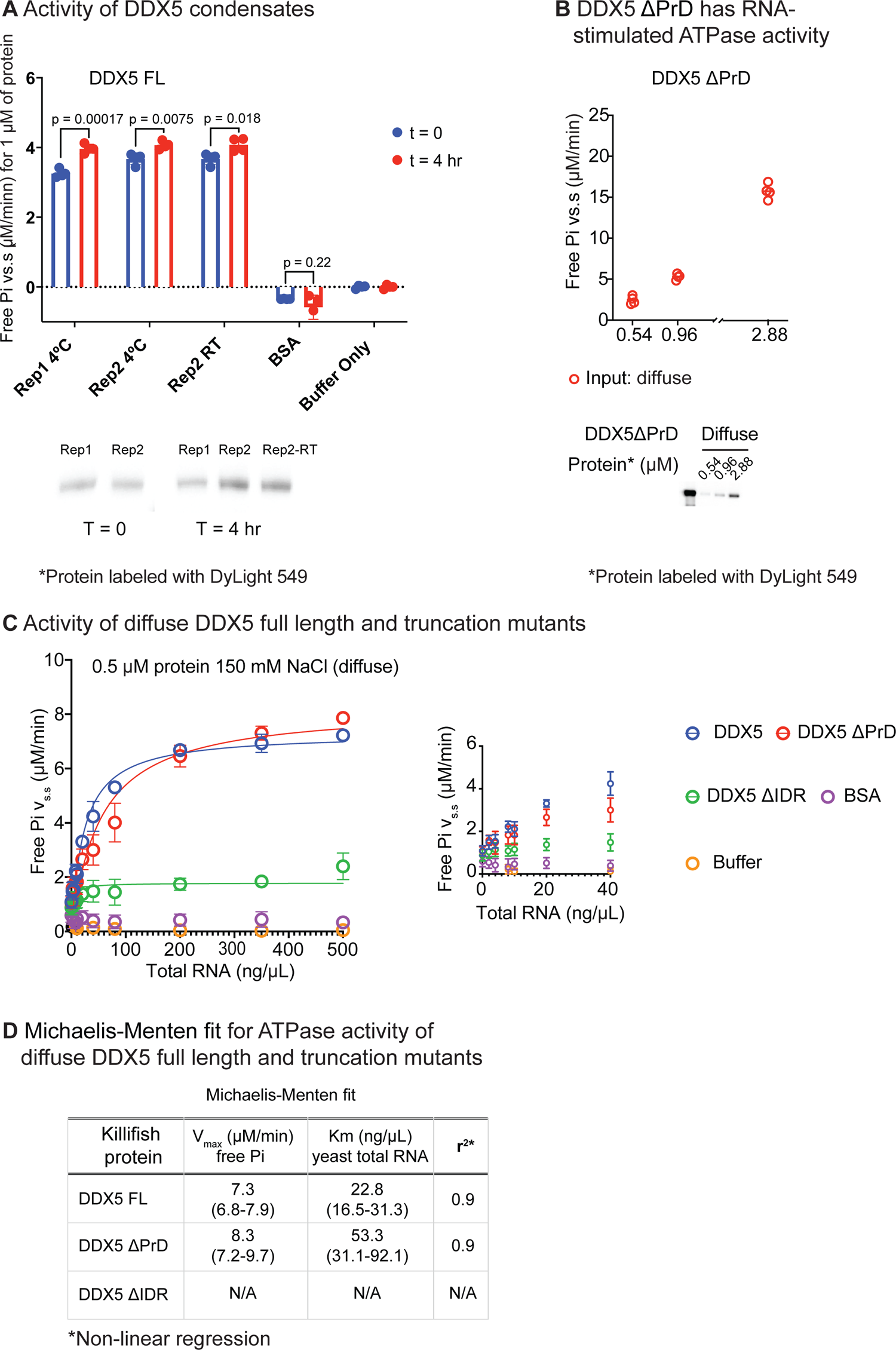
**(A)** ATPase assay results from an additional independent experiment (similar to Figure 6B). Killifish DDX5 protein in diffuse (red circles) or condensate (blue circles, above 2 μM) forms were used to catalyze ATP hydrolysis in the presence of yeast total RNA. The ATPase activity is measured as a function of free phosphate (Pi) release rate (μM/min), controlling for the same total DDX5 protein concentrations (µM). Values of 4 technical replicates are shown along with western blot to confirm a similar level of total DDX5 protein. **(B)** Killifish DDX5ΔPrD shows concentrations dependent ATPase activity in the presence of RNA (red circles). At these consecrations, the DDX5ΔPrD protein was in its diffuse form. ATPase activity is measured as a function of free phosphate (Pi) production rate (μM/min) at the indicated protein concentrations (µM). Values of 4 replicates from one representative experiment are shown along with SDS-PAGE of the reaction mixture. Two independent experiments were performed. **(C)** Left: ATPase activity for the killifish DDX5 full length (FL) and mutants (ΔPrD and ΔIDR). As negative controls, BSA (Bovine Serum Albumin) or buffer alone are used. Activity is measured under diffuse conditions (0.5 μM protein, 150 mM NaCl) as a function of free Pi production rate (μM/min) at the indicated RNA concentrations (ng/μL). Right: Inset displays activity at lower concentration with higher resolution. Mean +/- SEM from 3 independent experiments (3-4 replicates in each experiment) were shown. **(D)** Michaelis-Menten fit for the activity of DDX5 full length (FL) and mutants (ΔPrD and ΔIDR), as measured in (C). The Michaelis–Menten equation describes the rate of ATP hydrolysis by relating the rate of free Pi production (μM/min) catalyzed by DDX5 to the concentration of the RNA substrate (yeast total RNA) (ng/μL). The estimated maximum reaction rate (Vmax) and the Michaelis constant KM (a measure of the substrate-binding affinity) are provided.

## SUPPLEMENTAL TABLE LEGENDS

**Table S1.** Mass spectrometry dataset for the tissue lysate and protein aggregate fractions of brain samples in young, old, and old *TERT^Δ8/Δ8^* killifish – Related to Figure 1 and S1.

**Table S2.** Enriched gene sets from the Gene Set Enrichment Analysis (GSEA) in brain tissue lysate (TL), aggregate fraction (AGG), and aggregation propensity (PROP) from old compared to young animals – Related to Figure 1 and S1.

**Table S3.** Features of age-associated aggregates during brain aging in killifish – Related to Figure 1 and S1.

**Table S4.** Materials generated and used in this study.

